# Genomic and Immunogenomic Profiling of Extramedullary Acute Myeloid Leukemia Reveals Actionable Clonal Branching and Frequent Immune Editing

**DOI:** 10.1101/2025.04.18.649610

**Authors:** Clément Collignon, Tucker Hansen, Colin Hercus, Marianna B. Ruzinova, Gabrielle Roth Guepin, Caroline Bonmati, Marie Thérèse Rubio, Pierre Feugier, Mélanie Gaudfrin, Hervé Sartelet, Marion Divoux, Marc Muller, Sharon Heath, Geoffrey Uy, David Spencer, David Chen, Simona Pagliuca, Francesca Ferraro

## Abstract

Extramedullary acute myeloid leukemia (eAML) is a rare form of myeloid neoplasm characterized by leukemic infiltration outside the bone marrow (BM). Despite its prognostic significance, eAML is often underdiagnosed and poorly characterized at molecular level. We performed a comprehensive genomic and immunogenomic profiling on paired BM and extramedullary specimens from 26 eAML patients, alongside over 400 AML cases without extramedullary involvement and 97 healthy controls. Clonal branching from BM was observed in 38.5% of extramedullary sites, frequently involving actionable mutations in *FLT3*, *IDH2* and *NPM1* genes. Both compartments were enriched in *RAS* pathway mutations and class II HLA losses, suggesting active immunoediting mechanisms driving eAML development. Strikingly all relapsed cases acquired FLT3 aberrations, highlighting therapeutic opportunities. These findings underpin the need for improved detection and routine genomic profiling, including targeted sequencing of suspected extramedullary lesions.

## Introduction

Acute myeloid leukemia (AML) originates in the bone marrow (BM), and leukemic cells can escape this niche and infiltrate extramedullary sites, including the skin (*leukemia cutis*) and other tissues (*myeloid sarcoma*)—a phenomenon known as extramedullary acute myeloid leukemia (eAML)^1^. eAML can arise at various stages of disease progression, presenting at initial diagnosis, during relapse after chemotherapy, or following allogeneic hematopoietic cell transplantation (allo-HCT). Notably, it may occur in isolation, precede overt BM involvement, or develop alongside marrow disease^2–5^.

Although symptomatic eAML is reported in only ∼ 2% of AML cases^2,6–10^, the inherent limitations of current detection methods, including the inability to capture small or subclinical extramedullary lesions, combined with the absence of routine eAML evaluation in standard AML care, suggest that its true incidence may be higher. For instance, recent imaging-driven studies indicate a prevalence closer to 22%^11^. Furthermore, the recent European Leukemia Network (ELN) 2022 AML guidelines^7,8^ did not provide specific recommendations on prognosis and treatment for this distinct condition. To date, the prognostic significance of eAML remains controversial, with retrospective studies yielding conflicting results—some indicating a negative impact on overall survival, others finding no effect, and some suggesting that outcomes depend on established genetic risk factors^6,12–20^. A major reason for these limitations is the lack of direct characterization of eAML lesions, which are assumed to be an extension of BM disease rather than a distinct entity. This assumption has contributed to underdiagnosis of clinically inapparent lesions and limited molecular profiling in standard practice, leaving critical gaps in understanding the biology and prognostic implications of eAML. Therefore, more comprehensive genomic and immunogenomic characterizations are needed to refine risk assessment and to better define the risk factors for the development of extramedullary disease.

Previous genomic studies of eAML have primarily utilized targeted sequencing approaches with heterogeneous mutational panels^2,21–26^, which have provided valuable insights into cancer-associated genes but were not designed to comprehensively evaluate immune-related loci such as human leukocyte antigen (HLA) and Killer-cell immunoglobulin-like receptor (KIR) genes. While it is well-documented that the incidence of eAML significantly increases as a solitary relapse following allo-HCT^27,28^, the molecular mechanisms underlying this phenomenon remain poorly characterized. A well-established driver of post-transplant AML relapse is the downregulation^29,30^ or genomic loss^31^ of HLA, enabling leukemia cells to escape the graft versus leukemia (GvL) effect. However, apart from one case report^32^ reporting the loss of heterozygosity of HLA genes, no study has systematically investigated whether immune evasion mechanisms contribute to the pathogenesis of eAML.

In addition, comparative analyses of paired BM and eAML samples have been limited, making it challenging to fully characterize clonal evolution and the mechanisms underlying extramedullary dissemination. This study builds upon prior work by leveraging a more comprehensive genomic approach and paired sample design to advance our understanding of the molecular drivers of eAML and inform future strategies for clinical management.

Here we present a comprehensive genomic and immunogenomic analysis of paired eAML-BM specimens from a cohort of 26 patients, leveraging an integrative approach, illustrated in **Figure 1A**. Findings were contextualized through comparisons with two genomically characterized independent control cohorts: AML patients without extramedullary involvement from the BeatAML study^10,33^ and 97 healthy individuals^34^. Whole-exome and whole-genome sequencing were used to characterize point mutations, structural variants, and mutational signatures, along with in-depth immune profiling, including HLA and KIR genotyping.

**Figure 1.**
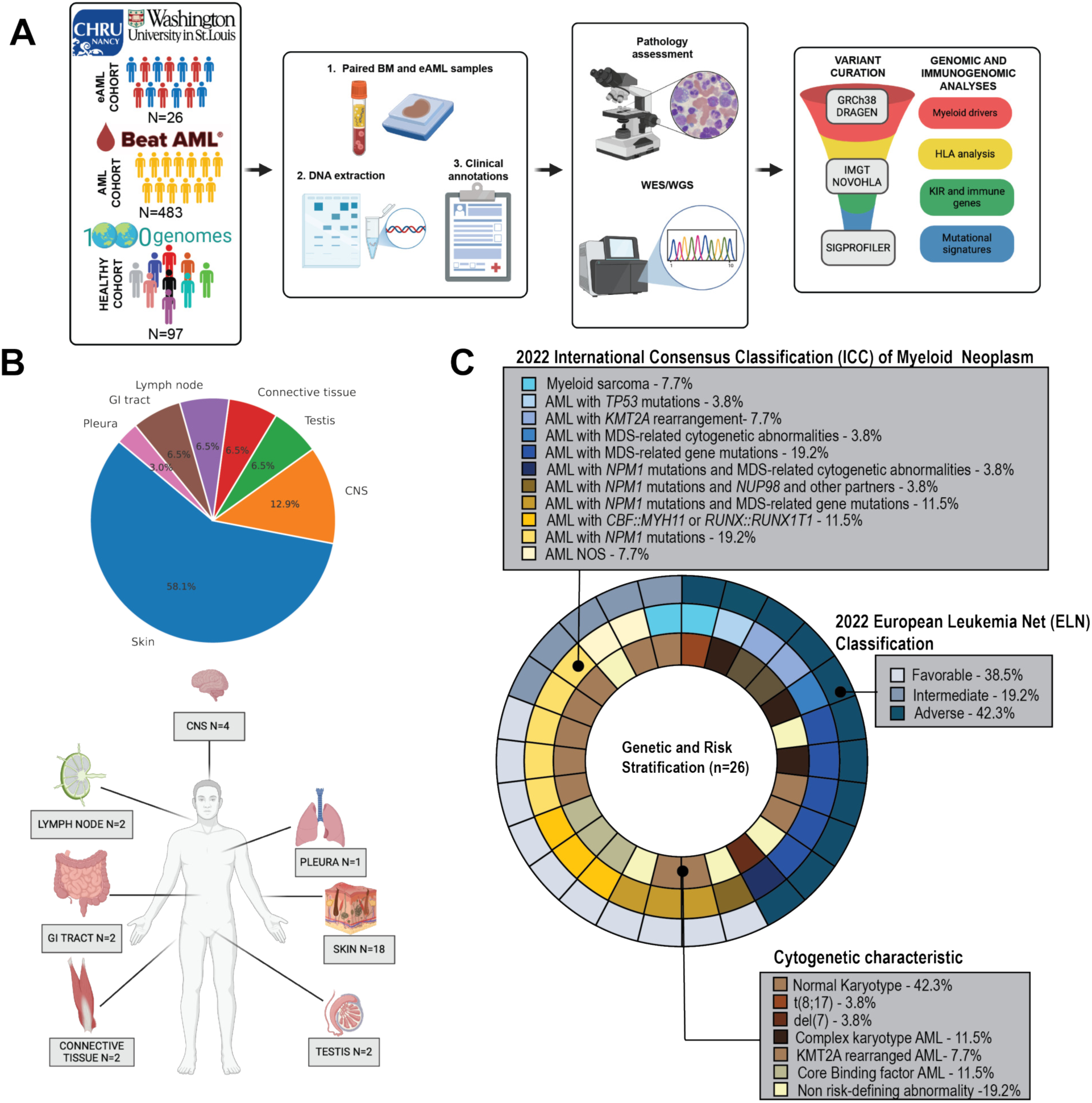
eAML study design and clinical characteristics. **(A) Study Design and Analytical Workflow.** This study analyzed paired bone marrow (BM) and extramedullary (eAML) samples from 26 patients treated at CHRU Nancy and Washington University in St. Louis. As a comparator, 483 AML cases without known extramedullary involvement from the Beat AML cohort were included. The workflow included: (1) collection of paired BM and eAML samples, (2) DNA extraction, and (3) comprehensive clinical annotation. Pathology assessment confirmed the diagnosis, followed by whole-exome sequencing (WES) and whole-genome sequencing (WGS) for high-resolution genetic and immunogenetic profiling. Variants were curated using GRCh38-aligned DRAGEN processing, IMGT-NOVOtyper for HLA typing, and SigProfiler for mutational signature analysis. The final analyses focused on (i) myeloid driver mutations, (ii) HLA profiling and immune escape mechanisms, (iii) KIR and immune gene assessments, and (iv) mutational signature characterization. **(B) Distribution of Extramedullary AML Sites.** The top panel shows a pie chart representing the frequency of eAML cases at different anatomical sites, while the bottom panel illustrates the anatomical distribution of disease involvement in the cohort. **(C) Genomic and Cytogenetic Landscape of eAML at Diagnosis.** The inner ring represents cytogenetic characteristics, the middle ring categorizes cases based on the 2022 International Consensus Classification (ICC), and the outer ring depicts risk stratification according to the 2022 European LeukemiaNet (ELN) criteria.

By examining the genetic and immune-related factors of paired eAML-BM samples, this study provides insights into clonal evolution, immune evasion strategies, and possible therapeutic targets, that could inform improved risk assessment and precision medicine approaches for patients presenting with, or at risk for developing, extramedullary disease.

## Results

### Patients and samples characteristics

The cohort consisted of 9 female (median age: 56 years, range: 25–65) and 17 male patients (median age: 64 years, range: 2–83). Twenty-three patients had de novo AML, two evolved from antecedent myelodysplastic syndrome and one case had therapy-related AML. The most common site of eAML involvement was the skin (58.1%), followed by the central nervous system (12.9%). Other affected sites included the testis, gastrointestinal tract, lymph nodes, lungs, and muscle (**Figure 1B**). Total white blood cell (WBC) count was 16.75 (Interquartile range—IQR 4-44.3) and median BM blast percentage was 53% (IQR 30-80%). Cases were further categorized based on the timing of eAML detection, occurring at initial diagnosis, at relapse following chemotherapy or at relapse after allogeneic transplant (**Table S1)**. BM concomitant to the extramedullary (paired) sample was available for 20 cases (**Figure S1A)**.

When eAML was detected at relapse, sequencing was also performed on the diagnostic BM sample. Additionally, for a subset of cases, multiple synchronous or metachronous eAML lesions were available for sequencing, providing insight into clonal evolution across different time points and anatomical sites (**Figure S1A, Table S2**).

According to the ELN 2022 classification^7^, at the time of AML diagnosis, 38.5% of patients had favorable-risk disease, 42.3% had adverse-risk disease, and 19.2% had intermediate-risk disease (**Figure 1C, Table S1**). At the time of analysis, no significant differences in overall survival based on ELN risk groups were detected (**Figure S1B**), highlighting the limitations of current risk stratification systems in accurately characterizing this AML subtype.

### Mutational analysis of paired BM and eAML samples reveals FLT3 enrichment and small inversions in eAML

First, we assessed the mutational status of thirty-four recurrently mutated genes (RMG) in AML (see **Methods**) in both eAML and BM specimens. Single nucleotide variant analysis revealed at least one pathogenic variant in RMG in 92% of the tested samples (**Figure 2A, Table S3**), including BM samples without morphological evidence of blasts. Notably, the only cases with no identifiable myeloid driver mutations in the BM occurred as relapses following allo-HCT (8_BMc, 27_BMc), in which *HLA* loss of heterozygosity (LOH) and deletions were detected. In the remaining BM samples, other AML drivers were implicated, such as *MYST3::CREBBP* translocation in case 1 and *IKZF1* mutation in case 24.

**Figure 2.**
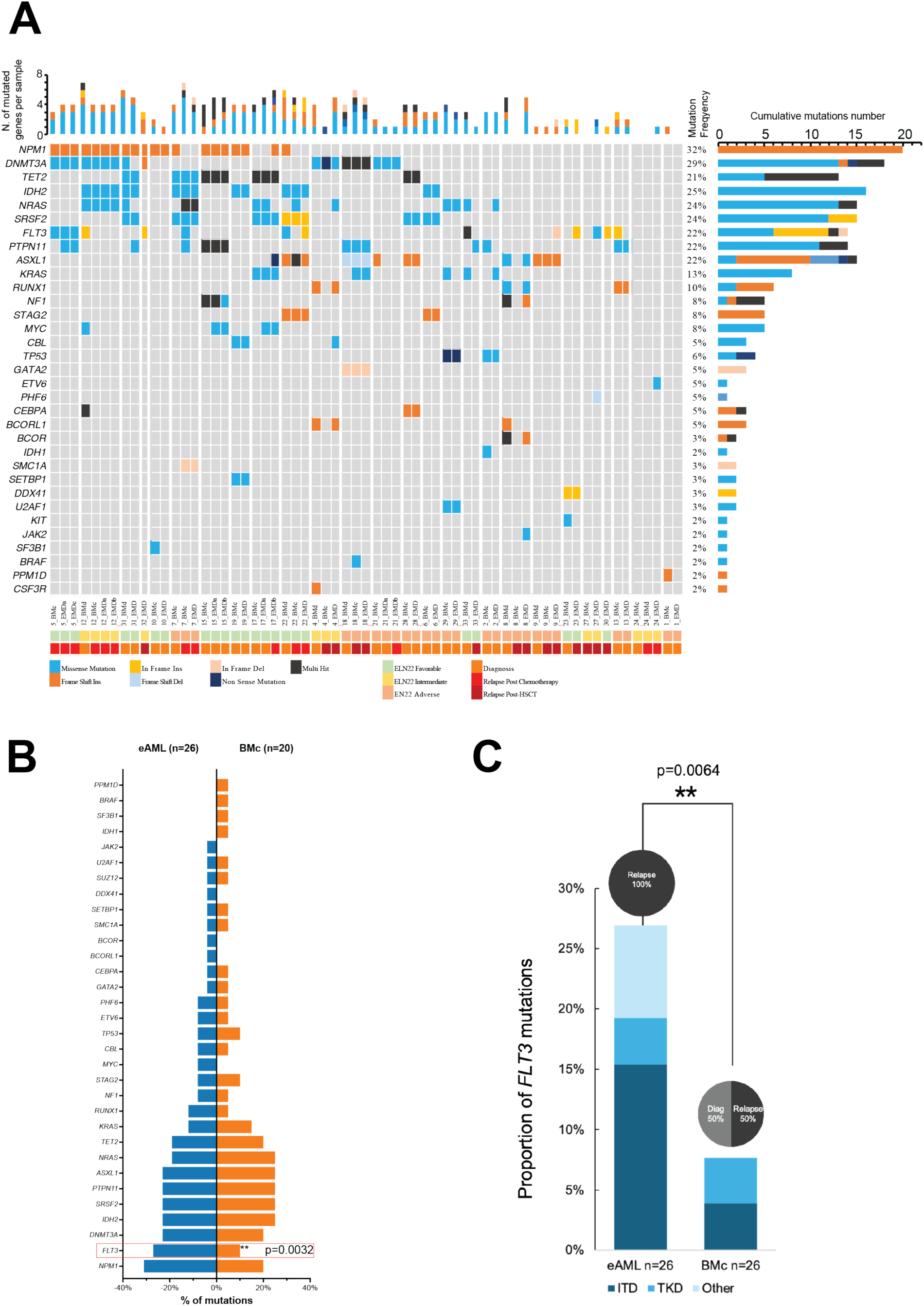
Mutational Landscape of Extramedullary Acute Myeloid Leukemia (eAML) Patients. **(A) Oncoplot of Recurrently Mutated Myeloid Genes in eAML Cohort.** Each column represents an individual patient, and each row corresponds to a specific gene. Mutation types are color-coded: missense mutations (blue), frameshift insertions (orange), inframe insertions (yellow), frameshift deletions (light blue), inframe deletions (light orange), nonsense mutations (dark blue), and multi-hit mutations (black). The annotation bar at the bottom indicates the clinical status of each patient: ELN22 favorable (light green), ELN22 intermediate (yellow), ELN22 adverse (light orange), diagnosis (orange), relapse post-chemotherapy (red), and relapse post-hematopoietic stem cell transplantation (dark red). The right panel displays the frequency of mutations per gene, with colors reflecting their distribution across different clinical conditions. **(B) Comparative Mutation Frequency Between eAML and Concomitant Bone Marrow Samples.** Bar plot illustrating the frequency of gene mutations in extramedullary AML samples (eAML, blue) compared to paired bone marrow samples (BMc, orange), highlighted is the enrichment of FLT3 mutations in eAML cases compared to paired BM samples (Fisher exact p=0.0032). **(C) FLT3 Mutation Frequency and Distribution in eAML and Bone Marrow Samples.** Stacked bar chart comparing the frequency and types of FLT3 mutations between extramedullary AML sites (eAML) and bone marrow (BM all) samples. FLT3 mutations, including internal tandem duplications (ITD) and tyrosine kinase domain (TKD) mutations, were more prevalent at relapse and significantly enriched in eAML sites compared to BM samples (Fisher exact p=0.0064).

The median number of mutations in myeloid-associated genes was comparable between BM and eAML samples (4 [IQR range 1.75-5.25] in BM vs. 4 [IQR range 2-5] in eAML, p=0.98, **Figure S2A**). Likewise, the median number of mutated myeloid genes was 3.5 [IQ range 1.75-4] in BM and 4 [range 2-5] in eAML (p=0.79 **, Figure S2B**). Similar results were observed comparing eAML samples and BM concomitant or at diagnosis. The most common mutations detected in eAML sites were *NPM1* and *FLT3* (31% and 27% respectively, **Figures 2A-B**), whereas in the paired BMs, the most frequent mutations were *IDH2*, *SRSF2*, *PTPN11*, *ASXL1* and *NRAS* (25%). The frequency of *FLT3* mutations was significantly different between eAML and paired BM samples (**Figure 2B**). The mutation rates were 25% in eAML and 10% in paired BM (Fisher exact p-value=0.0032) and the probability of having a FLT3 mutation in eAML cells was 3.3 times higher than in paired BM (Odds Ratio: 3.3, 95% CI: 1.51 – 7.32). Given the high prevalence of *FLT3* mutations in eAML specimens, we aimed to further investigate their significance. In total, we identified fifteen *FLT3* mutations: six in BM diagnosis (BMd) samples, two in paired, BM concomitant (BMc) samples, and seven in eAML samples (**Table S4**). Among the paired BMc/eAML samples, *FLT3* mutations were more prevalent in eAML samples. Specifically, BMc samples contained one *FLT3-ITD* and one *FLT3-TKD* mutations, whereas eAML samples harbored four *FLT3-ITD*, one *FLT3-TKD*, one *FLT3* I867S and one *FLT3 D600del* mutation. Notably, 2 *FLT3-ITD,* 1 *FLT3-TKD*, one *FLT3* I867S and one *FLT3*-D600del mutations were detected in the extramedullary samples and were absent in the paired BM sample and in the diagnosis BM. In contrast, in five cases, *FLT3* mutations originally present at diagnosis were no longer detectable in either the eAML or paired BM sample (**Figure S2C** and **Table S4**), highlighting significant clonal divergence between BM and extramedullary sites. Furthermore, all *FLT3* mutations in eAML were detected in relapse samples, whereas only one relapsed BM sample harbored a *FLT3* mutation (Fisher’s exact test, p = 0.0064; **Figure 2C**), highlighting the importance of comprehensive molecular profiling at relapse.

Next, we aimed to assess the degree of genomic instability associated with eAML by quantitatively and qualitatively analyzing structural variants (SVs) in both BM and extramedullary samples. While the overall proportion of SVs did not differ between paired samples (**Figure 3A-B**, **Figure S3A**), eAML exhibited an enrichment of small (<150 bp) SVs, particularly inversions (**Figure 3C, Figure S3B**). In contrast, large deletions (>500 kb) were predominantly observed in BM specimens at the time of eAML diagnosis (**Figure 3C**). Gene-level analysis revealed recurrent SVs affecting *CSF3R, NPM1, NRAS*, and *PTPN11*, with a higher prevalence in eAML samples (**Figure 3D**).

**Figure 3.**
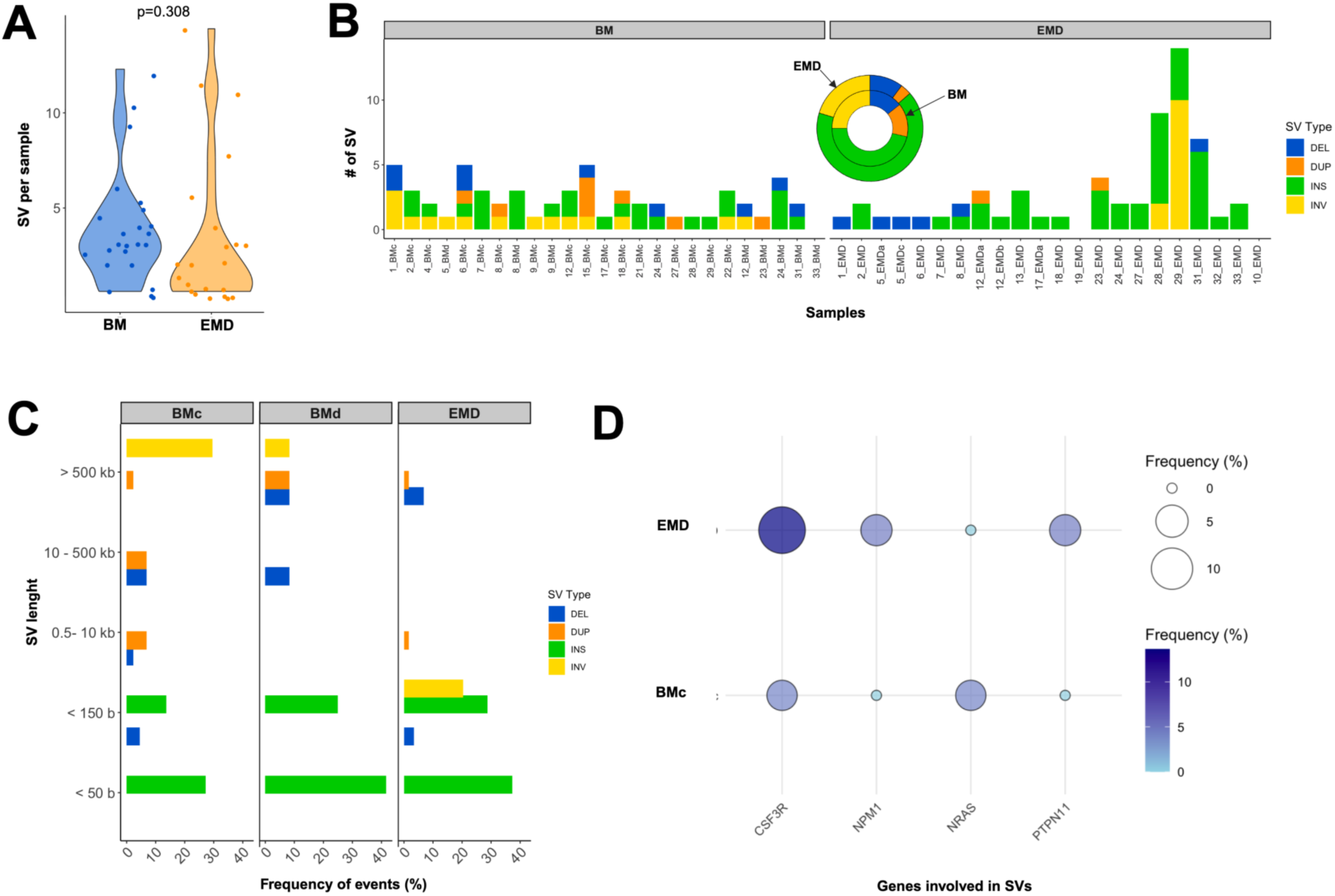
Structural Variant (SV) Landscape in Bone Marrow (BM) and Extramedullary Disease (EMD) (A) **Violin plot comparing the number of structural variants (SVs) per sample** between bone marrow (BM, in blue) and extramedullary disease (EMD, in orange), illustrating differences in SV burden (p=0.38, not significant). (B) **Bar plot displaying the number of SVs per sample across the cohort**, categorized by SV type: deletions (DEL, blue), duplications (DUP, orange), insertions (INS, green), and inversions (INV, yellow). The inset pie chart summarizes the overall distribution of SV types in BM and EMD. (C) **Frequency distribution of SVs by size** category across concomitant BM (BMc), BM at diagnosis (BMd), and EMD samples. SV types (DEL, DUP, INS, INV) are color-coded as in panel B. (D) **Bubble plot illustrating the frequency of SVs in myeloid-related genes**, comparing their distribution between BMc and EMD. The bubble size represents the proportion of samples harboring SVs in each gene, while color intensity indicates the relative frequency of these SVs within each group.

### RAS pathway mutations are more frequently detected in the bone marrow of patients with extramedullary disease compared to those without

To assess whether the mutational landscape differs between patients with extramedullary leukemia and those without, we compared the frequency of myeloid gene mutations detected in bone marrow samples (both at diagnosis and at the time of eAML localization) to a large dataset of de novo AML cases from the BeatAML trial^10^, after excluding cases associated with myeloid sarcoma (n=483, **Figure S4A**). This analysis revealed a significant enrichment of *RAS*/*MYC* pathway mutations in our cohort (p=0.0017; **Figure 4A, Figure S4A-B**), with *PTPN11* (p=0.0045), *NRAS* (p=0.38), *KRAS* (p=0.43), and *MYC* (p=0.024) being the most frequently enriched genes (**Figure 4B**). *RAS* pathway mutations were also enriched in the BM samples collected concurrently to the extramedullary disease (BMc, **Figure S4C-D**).

**Figure 4.**
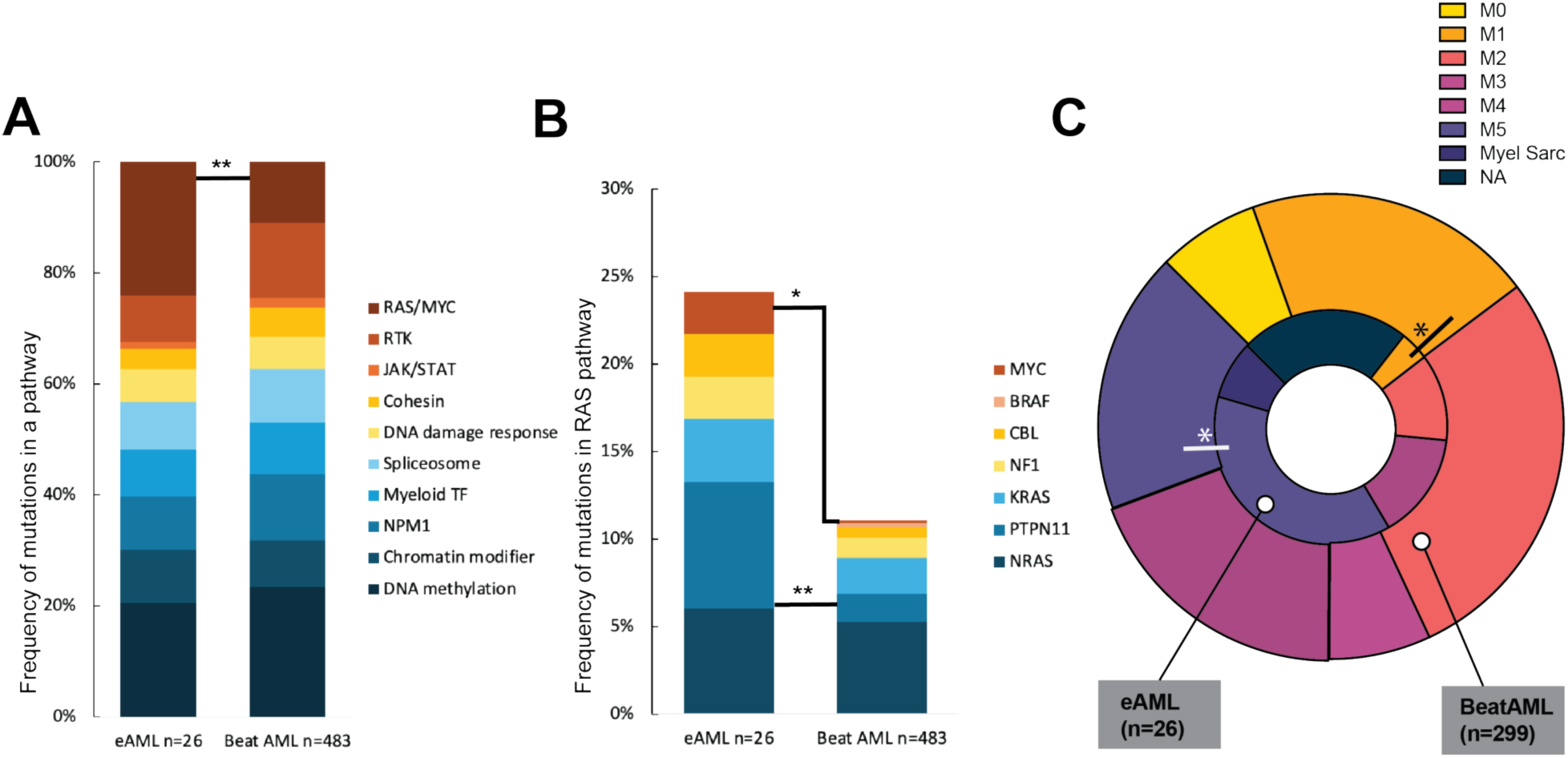
Comparative analysis of extramedullary AML (eAML) and the BeatAML cohort. **(A) Stacked bar plot illustrating the frequency of mutations across key functional categories in eAML (n=26) and BeatAML (n=483).** Categories include DNA methylation, chromatin modifiers, *NPM1*, myeloid transcription factors (TFs), spliceosome components, DNA damage response, cohesin, JAK/STAT signaling, receptor tyrosine kinases (RTKs), and the *RAS/MYC* pathway. **(B) Stacked bar plot comparing the prevalence of mutations in RAS pathway genes** (*NRAS*, *PTPN11*, *KRAS*, *NF1*, *CBL*, *BRAF*, and *MYC*) between eAML (n=26) and BeatAML (n=483). Significant differences are indicated: (*) p < 0.05; (**) p < 0.01. **(C) Donut chart comparing the distribution of French-American-British (FAB) AML subtypes between eAML (n=26) and BeatAML (n=299)**. Asterisks (*) indicate statistically significant differences in FAB subtype distribution between the two cohorts.

Given that *RAS* mutations are commonly associated with monocytic AML^35,36^, we next assessed the distribution of cases according to the FAB (French American British^37^) classification and compared it to *de novo* AML cases from the BeatAML trial. In our cohort, we observed a significant enrichment of M5 (monocytic) AML cases (p=0.018) and a significant depletion of M1 (AML with minimal differentiation) (p=0.038; **Figure 4C**), along with a trend toward depletion of M0 (undifferentiated AML) and M3 (acute promyelocytic leukemia), though not statistically significant (p = 0.23 and p = 0.39, respectively).

### BM and eAML sites often display divergent clonal evolution and mutational discordance

Next, we analyzed the clonal composition of eAML sites compared to BM samples by examining clonal proportions and normalized variant allele frequency (VAF), as detailed in the Methods section, and visualized the tumor evolution using fishplots^38^. This approach revealed distinct patterns of clonal evolution across paired samples. The most common pattern, observed in 65% of cases, involved discordant mutational profiles, where subclones undetectable in the BM were uniquely present in the eAML site (**Figures 5A-D**). In cases of isolated eAML, such as in relapses after allo-HCT, BM samples often contained only clonal hematopoiesis-related mutations, whereas the eAML site showed an ancestral clone that had acquired cooperating mutations (**Figures 5C-D**).The second most prevalent pattern was concordant mutational profiles with similar clonal expansion dynamics (35%, **Figure 5E**), suggesting shared evolutionary trajectories between the two compartments. However, in some cases, while the mutational profiles remained concordant, differences in clonal expansion between the BM and eAML site were observed (**Figure 5F**). Additionally, in some patients, progressive eAML lesions exhibited subclones derived from both the BM and the initial eAML site, highlighting a complex interplay of clonal evolution with contributions from multiple origins (**Figure 5G**). Notably, several eAML lesions harbored clinically actionable mutations, such as *FLT3*, *IDH2,* and *NPM1* (**Figures 5B-E, G**), as well as potentially targetable alterations, including mutations in the RAS pathway (**Figures 5A, D, E**). Due to the observed mutational discordance between BM and extramedullary eAML sites, we hypothesized that AML cells might be subjected to mutational stressors that may be different in the extramedullary sites compared to the BM. To determine whether AML cells residing in extramedullary sites displayed evidence of site-specific mutational signatures (i.e. UV-light mutational signatures for skin samples), we applied SigProfiler^39^ to extract mutational signatures from sequencing data, then compared the extracted mutational patterns against the COSMIC (catalogue of somatic mutation in cancer)^40,41^ reference database. Mutational signatures are characteristic patterns of single-base substitutions left behind by various mutagenic processes. These processes span endogenous cellular mechanisms (e.g. normal DNA replication errors or spontaneous deamination of 5-methylcytosine), environmental exposures (such as ultraviolet light or smoking-related DNA damage), and defective DNA repair pathways (for example, mutations arising from mismatch repair deficiency). Notably, despite the distinct microenvironmental pressures in extramedullary sites, we did not detect hallmark signatures associated with external carcinogenic exposures and BM and eAML sites displayed a largely overlapping mutational profile with no evidence of site-specific signatures (**Figure S6**). Specifically, the most prominent signatures were SBS1 and SBS5 (depicted in blue and orange, respectively). SBS1 is attributed to spontaneous deamination of methylated cytosine, while SBS5 represents a ubiquitous background mutational process linked to natural endogenous damage accumulated over time^42^. The third most prevalent signature is SBS54 (mustard-colored), which has been observed in various cancers but is not yet fully characterized –suggesting it may result from an as-yet unidentified mutagenic mechanism versus a sequencing artifact. Several other less frequent signatures are also detected occasionally and at lower levels, including SBS29, SBS31, SBS32, SBS37, SBS46, SBS54, and SBS87 (represented by various other colors in the bars). These minor signatures may correspond to specific exposures or DNA repair defects in certain samples. Overall, these findings highlight that the core mutational mark of AML is maintained across tissues.

**Figure 5.**
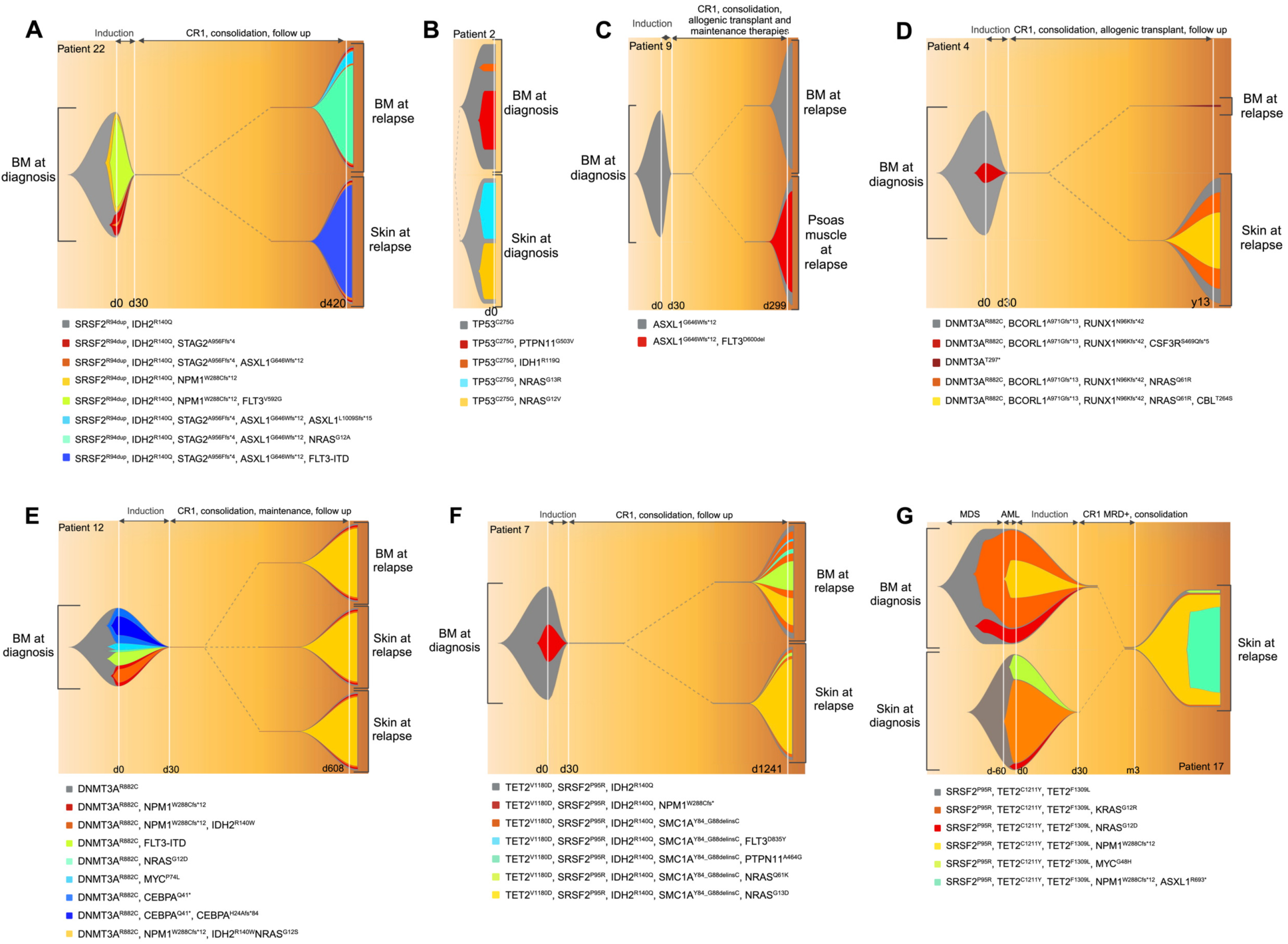
Clonal evolution patterns in bone marrow and extramedullary disease over the time. **(A-G) Fish plots illustrating distinct patterns of clonal evolution in patients with eAML**, depicting the dynamics of clonal architecture over time in bone marrow (BM) and extramedullary sites. Mutations are color-coded according to the legend. Panels (**A-C**) show cases of concomitant BM and eAML with discordant mutational profiles. These cases indicate divergent evolutionary trajectories and potential site-specific selective pressures driving leukemic progression. Panel (**D**) illustrates an isolated myeloid sarcoma relapse with no detectable BM involvement at relapse, demonstrating spatially restricted clonal evolution. Panel (E) represents a case of concordant mutational profiles in BM and eAML, with similar clonal expansion patterns, suggesting parallel evolution of leukemic clones across compartments. Panel (F) shows a case of concomitant BM and eAML with concordant mutational status but differential clonal expansion, indicating distinct evolutionary pressures shaping clonal dynamics in BM versus extramedullary compartments. Panel (**G**) presents progressive eAML lesions containing subclones derived from both BM and the in itial eAML site at diagnosis, highlighting intercompartmental clonal exchange and ongoing clonal adaptation.

### Immunogenetic landscape of eAML reveals immune escape through genetic HLA losses

Without a distinct mutational signature or unique drivers of extramedullary localization, along with the increased incidence of eAML relapses following allo-HCT, where immune editing is common^29–31,43–45^, we reasoned that genetic immune escape may play a role in establishing extramedullary disease. Therefore, we conducted a comprehensive assessment of both the germline (genotyping) and somatic (mutations) immunogenetic landscape of eAML. We first analyzed the allelic frequency of immune genes (**Table S5**) in our patient cohort and performed comparative assessments with healthy controls (**Table S6**). The goal of this analysis was to determine whether specific alleles of classical and non-classical *HLA* and *KIR* genes were preferentially associated with eAML. Allele enrichment analysis revealed no significant correlation between classical HLA alleles and the eAML phenotype. However, we identified a significant overrepresentation of specific activating and inhibitory KIR alleles, as well as distinct *MICA* and *MICB* alleles in eAML patients (**Figures 6A-C** and **Table S5**) compared to healthy controls. Given the differential distribution of *KIR* genes between eAML patients and healthy controls, we next investigated whether there was an imbalance in *KIR* ligand distribution within the *HLA-C* locus, which could contribute to NK cell dysfunction. *HLA-C* alleles function as KIR ligands and are classified into two groups based on their specificity: C1 (Asp80) and C2 (Lys80)^46^. Notably, *HLA-C2* homozygosity has been identified as a risk factor for post-allo-HCT leukemia relapse and has been linked to increased susceptibility to certain solid and hematologic malignancies^47–49^. To explore this potential association, we analyzed the genotypic distribution of the *HLA-C* locus and its correlation with the eAML phenotype. However, we found no significant differences in the frequency of C1/C2 homozygous or heterozygous genotypes between eAML patients and healthy donor controls (**Figure 6D**).

**Figure 6.**
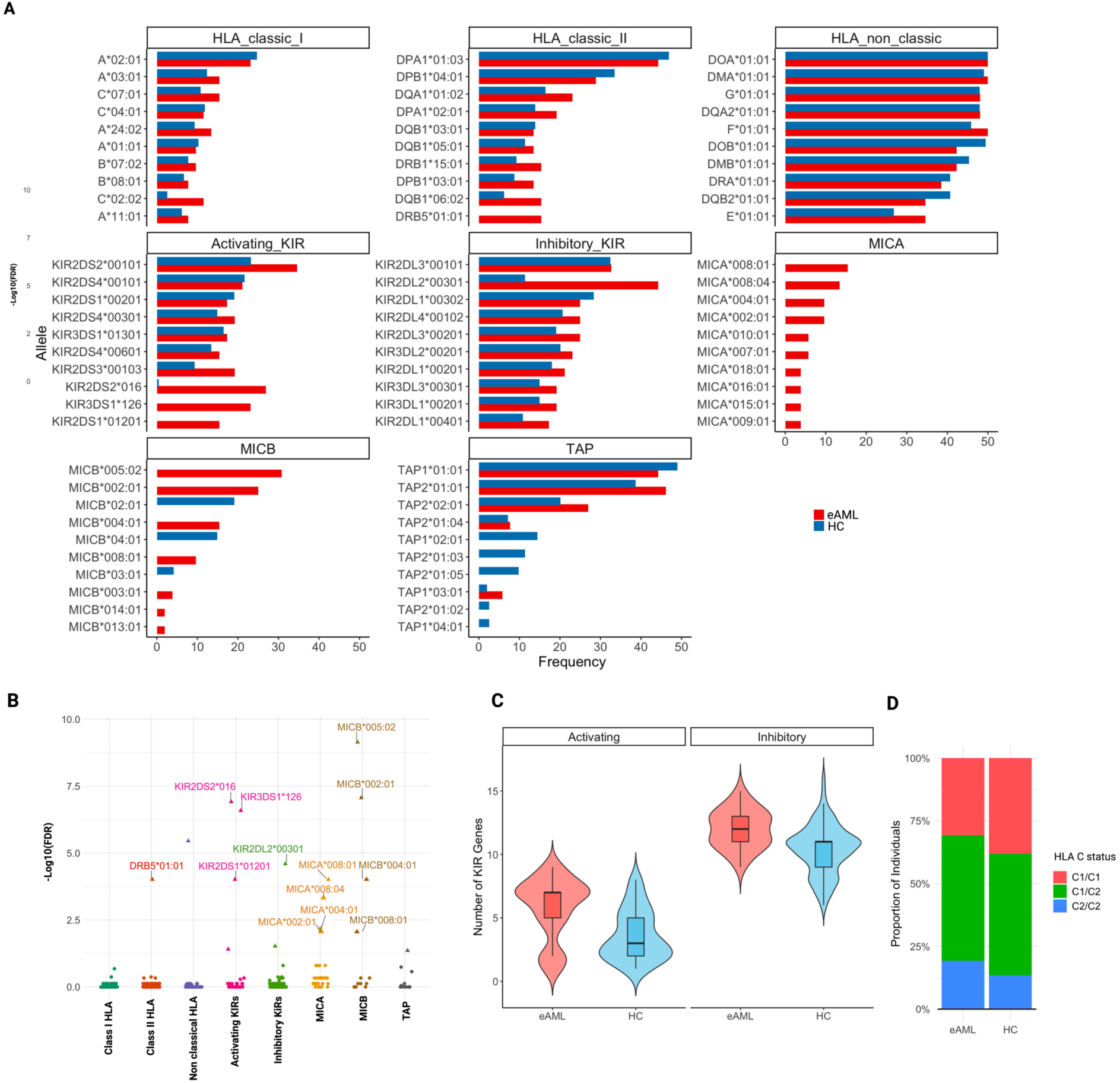
Immunogenetic profiling of extramedullary AML (eAML) compared to healthy controls (HC). **(A) Frequency distribution of HLA, KIR, MICA, MICB, and TAP alleles in eAML (red) and healthy controls (blue).** The panels depict allele frequencies for classical HLA class I, HLA class II, non-classical HLA, activating KIR, inhibitory KIR, MICA, MICB, and TAP genes. **(B) Enrichment analysis of immunogenetic alleles in eAML versus healthy controls.** The – log10(FDR) values for various alleles are shown, highlighting statistically significant overrepresentation in eAML, particularly within activating KIR alleles (KIR2DS2, KIR2DS1, KIR2DS5), inhibitory KIR alleles (KIR2DL3), and MICA/MICB alleles. **(C) Distribution of activating and inhibitory KIR gene counts in eAML and healthy controls.** Violin plots show the number of activating (red) and inhibitory (blue) KIR genes per individual in eAML patients and healthy controls. **(D) HLA-C status (KIR ligand classification) distribution in eAML and healthy controls**. Stacked bar plot showing the proportion of individuals with HLA-C genotypes categorized as C1/C1 (red), C1/C2 (green), or C2/C2 (blue).

Next, we explored whether somatic immune escape mechanisms involving classical and non-classical HLA genes could explain the extramedullary localization of AML cells in our cohort. Although no recurrent *HLA* missense mutations were identified in WES data, we observed frequent *HLA* losses, including deletions and LOH (**Figure 7A**). In total, we identified 35 samples harboring *HLA* losses, comprising 78 distinct events**—**63 deletions and 15 LOH—across major *HLA* class *I* and class *II* genes. When analyzing the incidence of *HLA* losses by site (BM vs. eAML) and the timing of eAML detection (diagnosis, relapse post-chemotherapy, or relapse post-allogeneic transplant), we found a significant enrichment of *HLA* losses in extramedullary sites (p=0.0216, **Figure 7B**), particularly at diagnosis and at relapse post-allogeneic transplant (Fisher’s exact p=0.027 and p=0.069 respectively**, Figure S7A**).

**Figure 7.**
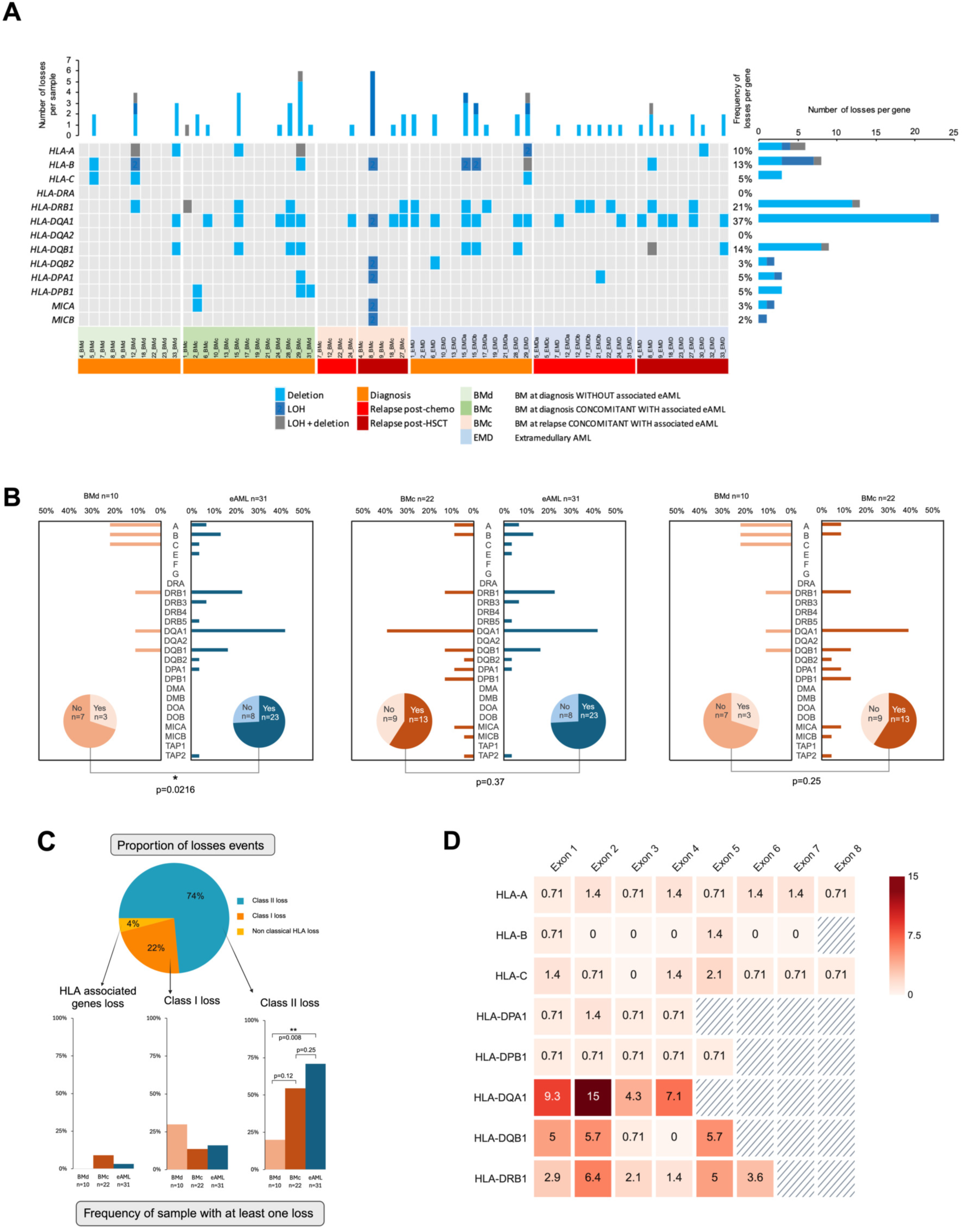
HLA Alterations in eAML Patients. **(A) Oncoplot illustrating the distribution of HLA losses across patient samples**. Each row represents an HLA gene, and each column corresponds to a patient. Blue shading differentiates types of HLA losses: deletions, loss of heterozygosity (LOH), or both. The bottom annotation bar categorizes patient groups by disease status: BM at diagnosis without eAML (BMd; light green), BM at diagnosis with concurrent eAML (BMc; green), BM at relapse with eAML (BMc; light orange), and extramedullary AML (EMD; light blue). The rightmost bar plot summarizes the percentage of samples affected by HLA losses for each gene. (B) **Comparison of HLA losses between BM and eAML samples**. Bar plots display the frequency of HLA losses in class I and class II genes across patient subgroups. Pie charts below each plot indicate the proportion of samples harboring at least one HLA loss. (C) **Proportion of HLA loss events across class I, class II, and non-classical HLA genes**. The top pie chart illustrates the relative distribution of HLA losses among these categories. Below, bar plots show the frequency of samples with at least one HLA loss, stratified by disease compartment and HLA gene class. (D) **HLA loss distribution across exons**. Heatmap depicting the frequency of deletions and LOH events across exons of different HLA genes. Darker shades indicate higher loss frequency, with exon 2 of HLA class II genes showing a notable concentration. Hatched areas mark exons with no detected losses.

Interestingly, in cases with concomitant BM and eAML involvement, leukemia cells from the BM exhibited *HLA* deletions or LOH at a slightly higher frequency than BM-derived cells from cases without synchronous eAML disease (59% vs. 30%, p=0.25), though this difference did not reach statistical significance. Notably, *HLA class II* genes were significantly more affected compared to *HLA class I* genes (74% vs. 22%, p<0.0001, **Figure 7C**). Specifically, while paired concomitant BM and eAML specimens showed no significant difference in class II losses (55% vs. 70%, p=0.25), these lesions were significantly more frequent in extramedullary sites compared to BM samples without concomitant extramedullary disease (70% vs. 20%, p=0.008).

When examining the distribution of HLA alterations across exons, we found that although the majority of losses and deletions occurred in exon 2 of class II genes—which encodes the antigen-presenting region of the HLA molecule—a consistent number of class I and II alleles were also altered outside this region, especially at sites involved in T cell interactions (e.g., CD8 or CD4 coreceptor contact sites) and in regions affecting the leader sequence in exon 1, which is critical for the proper folding, transport, and function of the HLA protein (**Figure 7D**).

To further explore the interplay between *HLA* alterations and recurrent myeloid gene mutations, we analyzed the association between *HLA* status and the mutational landscape in eAML and BM samples. Interestingly, despite the overall enrichment of *HLA* losses in eAML samples, cases harboring *NPM1*^50,51^ and *FLT3*^52^ mutations, both of which have been associated with epigenetic downregulation of MHC class II, tended to retain an unaltered HLA status within extramedullary sites (**Figure S7B**).

## Discussion

The mechanisms underlying leukemia dissemination to extramedullary sites remain poorly understood, and whether infiltration outside the BM is unequivocally related to intrinsic blast abnormalities or to an extrinsic failure of immune surveillance remains an outstanding question.

Through the application of whole-exome and whole-genome sequencing, this study provides novel insights into the genetic and immunogenetic landscape of eAML. By leveraging these approaches, we were able to characterize aberrations in immune-related genes and uncover distinct genetic features of leukemia cells residing in extramedullary sites compared to their BM counterparts.

Our findings highlight that eAML frequently exhibits clonal divergence from paired BM samples, with 65% of cases displaying discordant mutations. This suggests that leukemia cells in extramedullary sites undergo distinct selective pressures, likely related to the different spectrum of immune surveillance mechanisms, leading to the expansion of specific subclones. In some instances, eAML even appears to evolve independently from a pre-leukemic population rather than originating from an overt BM leukemic clone. Our case (Figure 5D) in which post-transplant relapse was restricted to eAML while BM harbored only myeloid (non-driver) mutations, suggests that eAML may evolve independently from a pre-leukemic population in extramedullary sites. Notably, the absence of a distinct mutational signature in eAML suggests that organ-specific mutagens are unlikely to drive this divergence, implicating alternative, immune-mediated mechanisms in the selection of extramedullary clones.

Importantly, we identified a high prevalence of mutations in genes such as *FLT3*, *RAS*, *NPM1*, and *IDH2* in eAML. These genes are considered actionable because they are associated with targeted therapies, either already approved or currently under investigation, and their identification can directly inform treatment selection. These findings underscore the importance of sequencing extramedullary lesions to uncover therapeutically relevant alterations that may not be detected through bone marrow analysis alone. *FLT3* mutations, especially *FLT3-ITD*, were enriched in eAML compared to BM, suggesting a potential role in the survival and expansion of leukemia cells in extramedullary sites. The presence of significant clonal divergence between BM and eAML further underscores the dynamic nature of leukemia evolution, with extramedullary organs potentially serving as reservoirs for subclonal populations that contribute to relapse.

We also observed a significant enrichment of *RAS* pathway mutations in both BM and eAML samples from cases with eAML compared to those without. This aligns with a previous meta-analysis of BM samples from eAML patients^25^ and is consistent with the known role of RAS mutations in promoting monocytic differentiation^35^. In our cohort, eAML cases were significantly enriched for M5 (monocytic) AML and depleted of M1 (minimally differentiated) AML, further supporting the association between *RAS* mutations, monocytic differentiation, and extramedullary dissemination.

Interestingly, germline *RAS* pathway mutations in genes such as *PTPN11*, *SOS1*, *KRAS*, *NRAS*, *RAF1*, *BRAF*, and *CBL* are implicated in a group of congenital disorders known as RASopathies, which include Noonan syndrome, Costello syndrome, cardiofaciocutaneous syndrome, juvenile myelomonocytic leukemia (JMML), and Neurofibromatosis type 1^53^. Notably, JMML is associated with an increased risk of myeloid malignancies^54^ and leukemia cutis^55^, reinforcing the link between *RAS* mutations and extramedullary leukemia. Moreover, *RAS* mutations have been implicated in the acquisition of invasive phenotypes^56,57^, suggesting that these genetic alterations may promote extramedullary dissemination and establishment of AML cells, thereby increasing the risk of eAML development. Given these findings, the presence of *RAS* mutations in BM samples should prompt careful evaluation for extramedullary disease, as these cases may be at higher risk for eAML progression.

Given that N-RAS and K-RAS inhibitors are in clinical trials for multiple cancers,^58–61^ and MEK inhibitors, which target downstream components of the *RAS* pathway, are already FDA-approved for melanoma^62,63^, these agents warrant further evaluation in the context of eAML. Combining RAS pathway inhibitors with standard chemotherapy may reduce the risk of relapse and mitigate extramedullary dissemination Beyond mutational analysis, our study also revealed frequent *HLA* deletions and LOH in eAML, particularly in *HLA class II* genes, both in antigen-presenting regions and other sites important for T cell interactions and peptide expression. These losses were significantly more common in extramedullary sites and, intriguingly, were observed not only at relapse after allogeneic transplantation but also at diagnosis, when compared to bone marrow samples. Interestingly, in cases with concomitant BM and eAML, BM-derived leukemia cells also exhibited higher frequencies of *HLA* deletions or LOH compared to cases without eAML, suggesting that immune escape mechanisms are already present in BM before clinically evident extramedullary dissemination, and they may be selected for in extramedullary sites.

The predominance of *HLA* class II exon 2 losses—which encode the antigen-presenting region— suggests a distinct pattern of immunoediting, potentially driven by specific antigens presented on the surface of AML cells. Moreover, the presence of mutations in regions that interact with CD4 (exon 3 for class II) or CD8 coreceptors (exon 4 for class I) may reflect T cell–mediated selective pressure. Notably, the lower frequency of *HLA* losses in *NPM1*-mutated AML is consistent with the known association between *NPM1* mutations and epigenetic downregulation of HLA class II expression^50,64^, suggesting that these two mechanisms may converge to promote immune evasion. These findings indicate that *HLA* loss is a key immune escape mechanism in eAML, particularly affecting *HLA* class II genes, which may contribute to leukemia persistence in extramedullary tissues. Indeed, when considering both genetic immunoediting events (e.g., deletions or loss of heterozygosity) and epigenetic events (e.g., mutations known to downregulate antigen presentation machinery), major histocompatibility antigen (MHC) deregulation was observed in 89% of the samples (**Figure S7C**), reinforcing the notion that distinct evolutionary pressures may contribute to immune evasion and extramedullary dissemination in AML.

Routine BM aspirate analysis could help identify eAML risk factors, with *RAS* pathway mutations and *HLA* deletion/LOH serving as biomarkers to guide proactive screening and clinical decision-making for extramedullary involvement.

In conclusion, our study underscores the importance of tissue sampling and sequencing of suspected eAML lesions to identify potential actionable mutations and guide treatment decisions. The detection of *RAS* pathway mutations and *HLA* losses in BM samples could serve as biomarkers for eAML risk, prompting closer surveillance, proactive imaging, and consideration for targeted therapies. Finally, the identification of clonal divergence in eAML sites supports the integration of eAML into measurable residual disease (MRD) monitoring strategies to decrease the risk of disease relapse and improve patient outcomes. Clonal divergence underscores the importance of biopsying and sequencing suspected eAML lesions when clinically feasible. To improve long-term remission, clinical trials should prioritize targeting eAML-specific mutations and incorporating eAML into measurable residual disease monitoring strategies.

## Methods

### Case selection

Cases were retrospectively identified from 2004 to 2023 through electronic medical records from two institutions: Centre Hospitalier Régional Universitaire (CHRU) de Nancy, Nancy, France, and Barnes-Jewish Hospital at Washington University School of Medicine, Saint-Louis, MO, USA (WashU). Selection criteria included the availability of formalin-fixed paraffin-embedded (FFPE) eAML tissue samples and concomitant cryopreserved BM DNA. Informed consent for the use of the residual biological specimens from the clinical sampling for sequencing purposes was obtained under Institutional Review Board (IRB)-approved protocols at both institutions (IRB Nancy n° 2024PI205-290, IRB WashU 201011766). See “Patients’ clinical summaries” in Supplementary appendix.

A control cohort of 97 healthy individuals (approximately a 1:4 case-to-control ratio) was retrieved from the 1000 Genomes Project^34^ to match the sex and race distribution of the cases and enable comparisons of the frequency distribution of classical and non-classical HLA genes, class I-like genes, and KIR genes.

A “disease” control cohort was retrieved from the Beat AML project^10^ which excluded subjects with known extramedullary disease.

### DNA extraction and sequencing

WES was performed at the McDonnell Genome Institute (MGI) sequencing facility at WashU. DNA was extracted from FFPE for eAML samples and BM biopsies, and BM aspirates using commercially available kits (Qiagen QIAamp DNA FFPE Tissue Kit and Qiagen DNeasy Blood & Tissue Kit) following manufacturer protocols. Genomic DNA (100–250 ng) was fragmented using a Covaris LE220 to achieve an approximate fragment size of 200–375 bp.

Libraries were prepared using the KAPA Hyper Prep Kit (KAPA Biosystems, Cat #7962363001) on a Perkin Elmer SciClone G3 NGS (96-well configuration) automated workstation. Libraries were pooled at equimolar ratios, generating up to 5 µg per pool, and hybridized using the xGen Exome Research Panel v2.0 reagents (IDT Technologies). Custom Illumina adapters with 10-base dual unique indexes were included. Pooled libraries were sequenced to generate paired-end reads of 151 bases on an Illumina NovaSeq X Plus instrument.

The average sequencing coverage across samples was 246×, with an average mapping rate of 99.07%. Base calling and demultiplexing were performed using the BCL Convert utility, either onboard or through a standalone DRAGEN processor, yielding sample-specific FASTQ files.

### Bioinformatics Workflow

Sequencing reads were analyzed on a DRAGEN Bio-IT processor running software version 4.2.4 in tumor-only mode, with and without germline tagging enabled. Reads were aligned to the GRCh38 reference genome, generating alignments in CRAM format. Small variants, including single nucleotide variants (SNVs) and indels, as well as copy number (CNVs) and structural variants (SCNVs), were called and vcf files generated.

### Post-Processing Filters variant analysis

SNVs variants were annotated for functional impact using ANNOVAR (version 2020-06-08) and the Variant Effect Predictor (VEP, version112, May 2024) while AnnotSV (version 3.4.1, 2024-05-03) was used for structural variants annotation. Post-variant calling filters included: variants with a minimum read depth of 10 and minimum alternate allele counts of 3 for genes recurrently mutated in AML (myeloid genes: *ASXL1*, *BCOR*, *BCORL1*, *BRAF*, *CALR*, *CBL*, *CEBPA*, *CHEK2*, *CSF3R*, *CUX1*, *DDX41*, *DNMT3A*, *ETV6*, *EZH2*, *FLT3*, *GATA1*, *GATA2*, *IDH1*, *IDH2*, *JAK2*, *KIT*, *KRAS*, *MYC*, *NF1*, *NOTCH1*, *NPM1*, *NRAS*, *PHF6*, *PIGA*, *PPM1D*, PTPN11, *RAD21*, *RUNX1*, *SETBP1*, *SF3B1*, *SMC1A*, *SMC3*, *SRSF2*, STAG2, *SUZ12*, *TET2*, *TP53*, *U2AF1*, *UBA1*, *WT1*, *ZRSR2*), and of 5 for other, non-myeloid genes. All variants identified through the workflow were visually reviewed using Integrative Genomics Viewer^65^ (IGV) to manually confirm each of the calls. Synonymous variants were excluded from downstream analyses.

### Handling of Germline Variants

Potential germline SNVs were flagged during the germline-tagging process. Variants with high population allele frequencies (>0.01%) in public databases (e.g., gnomAD, Cosmic, ExAC, 1000genome) were excluded unless strong literature evidence suggested pathogenicity in AML contexts. eAML variants were cross-referenced against patient-matched clinical sequencing. For structural variant analysis, insertions, duplications, deletions, were considered if classified by VEP as somatic and had a somatic score>30. All variants underwent careful manual curation to ensure that only highly confident pathogenic and likely pathogenic somatic calls would be used for the final analyses.

### Clonal comparison and clonal evolution analysis

To assess the clonal composition of eAML sites relative to bone marrow samples, we implemented a normalization strategy to account for differences in leukemia cell burden across tissues. BM aspirates exhibited substantial variability in leukemia cell content depending on the timing of collection, while eAML biopsies frequently contained varying proportions of non-tumor tissue.

To facilitate meaningful comparisons of VAF between these tissues, we applied the following normalization formula:

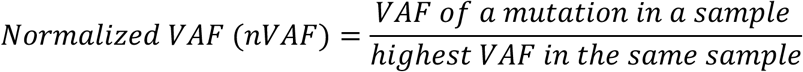

This approach adjusts each mutation’s VAF relative to the highest VAF mutation in the same sample, enabling a standardized assessment of clonal proportions independent of absolute VAF values. By normalizing VAFs in this manner, we ensured robust and reliable comparisons of clonal composition between eAML and bone marrow samples, even under conditions of variable tumor content.

### Mutational Signatures Analysis

Mutational signature analysis was performed using SigProfilerAssignment^39^, leveraging de novo extraction of mutational patterns from sequencing data. Identified signatures were compared against established COSMIC^41^ mutational signatures (v3.4) to characterize mutational processes specific to extramedullary eAML cells or identifying potential environmental or endogenous factors driving itsAML cell genetic evolution in extramedullary sites.

### Immunogenomic analyses

HLA and KIR genotyping from WES samples was performed using the NovoHLA pipeline, which was developed and validated in our prior work^31,66,67^ and is currently undergoing the licensing process by Novocraft Technology (https://www.novocraft.com/). This tool enhances the human reference genome (GRCh38.p13) with alternate scaffolds from IMGT (v3.57) and IPDKIR (v2.13) databases, allowing precise allele typing. Reads mapped to classical and non-classical HLA, MICA, MICB, TAP1, TAP2 and KIR genes, as well as unmapped reads, were extracted from BAM files based on GRCh38 coordinates and processed into a new BAM file.

This file was subsequently converted back to FASTQ format for allele typing. The typing process proceeds in three steps:

1) Initial Mapping: Reads are aligned to the reference genome with alternate allele scaffolds, and allele pairs are ranked based on coverage and alignment quality.
2) Refined Pileup: Variant calling and allele pair scoring are performed for candidates identified in the first pass, further refining pair rankings.
3) Consensus Matching: Variants in exons are compared with consensus coding sequences (CDS) to identify potential matches with alleles annotated only at the nucleotide level in the IMGT database.

Finally, the most likely allele pair for each gene is determined, and results are summarized in output files containing alignment details, variant scores, and coverage statistics.

HLA analysis was performed as previously described^31,66,67^. Briefly, For classical and non-classical HLA genes allelic loss was determined by calculating the number of reads covering each heterozygous allele within a given locus, using the following formula:

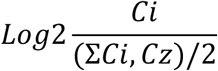

With Ci and Cz represent the read coverage for each allele within the same locus. For structurally similar alleles, an adjustment for sequence variation, taking into account the number of polymorphisms between two alleles, was applied, as directly computed by the NovoHLA pipeline. All Log2 ratios below –1.0 were classified as confident allelic losses, based on a prior internal validation study on other WES sample cohorts.

A LOH event was identified when heterozygous allele calls disappeared in the sample genotyping. Clinical allelic typing at the 2-digit resolution level was available for all patients and used as a reference.

For the healthy control cohort, since only 2-field genotyping data limited to HLA-A, HLA-B, HLA-C, HLA-DRB1, and HLA-DQB1 alleles were available, we utilized CRAM files from WGS available at https://ftp.1000genomes.ebi.ac.uk/ to extract the full HLA and KIR regions with the same pipeline.

## KIR-HLA interaction genotype analysis

The activity of each KIR gene and relative interactions with Class I HLA alleles were analyzed based on previous studies.^46,47–49^

KIR genes were categorized as either inhibitory (KIR2DL1, KIR2DL2, KIR2DL3, KIR2DL4, KIR2DL5A, KIR2DL5B, KIR3DL1, KIR3DL2, KIR3DL3) or activating (KIR2DS1, KIR2DS2, KIR2DS3, KIR2DS4, KIR2DS5, KIR3DS1), with genotyping performed using the NovoHLA pipeline as described earlier. KIR ligands were identified based on HLA-C typing, with C1 and C2 ligands assigned for interactions with KIR2DL1, KIR2DL2, and KIR2DS1^46^. The homozygous or heterozygous state of KIR ligands was determined according to their respective ligand class.

## Statistical analysis

Mean and 95% confidence intervals (CI) or median, interquartile ranges (IQR) were used where appropriate. Frequency and distribution of categorical variables were expressed as percentage. Descriptive statistics of continuous variables were reported as mean ± standard deviation (SD) for normally distributed data or median with interquartile range (IQR) for non-normally distributed data. For all relevant comparisons, after testing for normal distribution, comparative analysis between two groups were performed by two-sided paired or unpaired Student’s t-tests at 95% CI. In cases of not normally distributed data, the Wilcoxon matched pairs signed rank test at 95% CI was used. Fisher’s exact test or Chi-square were applied for independent group comparisons. In cases of testing more than two groups, a one-way ANOVA test was used.

Estimated effect size (odds ratio) was calculated when appropriate for group comparison (i.e. allele enrichment analysis). Multiple testing correction was performed using the Benjamini-Hochberg procedure.

Survival analysis was conducted using Kaplan-Meier curves, and differences in survival between groups were evaluated with the log-rank test. Overall survival (OS) was defined as the time from AML diagnosis to the last follow-up or death by any cause.

All statistical analyses and data visualizations were performed using GraphPad Prism (version 10.4.1), Microsoft Excel 365, and R software (version 4.2.0). The R environment used for the analyses in this manuscript included the following packages: ggplot2, ggsci, RColorBrewer, ggrepel, ggforce, gridExtra, Cairo, dplyr, tidyverse, reshape2, data.table, janitor, survminer, and maftools. Figures were assembled using BioRender and Adobe Illustrator (version 29.2.1).

## Conflict-of-interest disclosure

The Authors declare no competing interests.

## Data availability

All the data that support the findings of this study are available within the Article and Supplementary Files.

Source data are also provided in this manuscript. Sequencing data have been deposited under controlled access at the following repository 10.6084/m9.figshare.28661279. For any request, please contact the corresponding authors.

## Acknowledgements

We thank Agata Gruszczynska for help with the initial fastq files processing. We are grateful to the McDonnell Genome Institute at Washington University School of Medicine for their sequencing and genomic expertise. The Center is partially supported by NCI Cancer Center Support Grant #P30 CA91842 to the Siteman Cancer Center. We would like to thank Siteman Cancer Center’s scientific editor, Megan Noonan, PhD, for her expertise, meticulous attention to detail, and insightful suggestions, which enhanced the clarity and quality of this work. This work was supported by Leukemia Research Foundation grant to FF and by Fondation ARC pour la Recherche sur le Cancer, MDS Foundation Tito Bastianello Award, Force Hemato to S.P. Sample collection for Washington University is supported by Specialized Program of Research Excellence in Acute Myeloid Leukemia grant (P50 CA171963). We thank all the patients and their families who agreed to participate in this research. Notably, data collection, study design, and analysis were conducted as part of CC’s Master of Science program, with academic support from the Faculty of Medicine at the University of Lorraine, within the framework of research and training programs for young medical students.

## Authorship contributions

CC, SP and FF designed the study, collected, analyzed and interpreted the data, performed the bioinformatic and statistical analyses, and wrote the manuscript. SP and CH performed immune gene analysis and visualization. CH developed NovoHLA and the methodology for the calculation of copy number variation. GRG, MTR, CB, PF, MR, SH, GU participated in patient recruitment and management. MG, HS, MM, MD participated in sample recruitment. DS helped in data collection and interpretation and supervised genomic experiments. DC and TH participated in study conception, sample and data collection, data interpretation, gave important intellectual inputs, and edited the manuscript. All authors participated in the analysis interpretation and critical manuscript revision.

## Notes

https://doi.org/10.6084/m9.figshare.28661279.v1

### Competing Interest Statement

The authors have declared no competing interest.

## REFERENCES

1. Shallis, R.M., et al. Myeloid sarcoma, chloroma, or extramedullary acute myeloid leukemia tumor: A tale of misnomers, controversy and the unresolved. Blood Rev 47, 100773 (2021).

2. Abbas, H.A., et al. Clinical and molecular characterization of myeloid sarcoma without medullary leukemia. Leuk Lymphoma 62, 3402–3410 (2021).

3. Avni, B. & Koren-Michowitz, M. Myeloid sarcoma: current approach and therapeutic options. Ther Adv Hematol 2, 309–316 (2011).

4. Bakst, R.L., Tallman, M.S., Douer, D. & Yahalom, J. How I treat extramedullary acute myeloid leukemia. Blood 118, 3785–3793 (2011).

5. Wilson, C.S. & Medeiros, L.J. Extramedullary Manifestations of Myeloid Neoplasms. Am J Clin Pathol 144, 219–239 (2015).

6. Al-Khateeb, H., Badheeb, A., Haddad, H., Marei, L. & Abbasi, S. Myeloid sarcoma: clinicopathologic, cytogenetic, and outcome analysis of 21 adult patients. Leuk Res Treatment 2011, 523168 (2011).

7. Dohner, H., et al. Diagnosis and management of AML in adults: 2022 recommendations from an international expert panel on behalf of the ELN. Blood 140, 1345–1377 (2022).

8. Khoury, J.D., et al. The 5th edition of the World Health Organization Classification of Haematolymphoid Tumours: Myeloid and Histiocytic/Dendritic Neoplasms. Leukemia 36, 1703–1719 (2022).

9. Shahin, O.A. & Ravandi, F. Myeloid sarcoma. Curr Opin Hematol 27, 88–94 (2020).

10. Tyner, J.W., et al. Functional genomic landscape of acute myeloid leukaemia. Nature 562, 526–531 (2018).

11. Stolzel, F., et al. The prevalence of extramedullary acute myeloid leukemia detected by (18)FDG-PET/CT: final results from the prospective PETAML trial. Haematologica 105, 1552–1558 (2020).

12. Avni, B., et al. Clinical implications of acute myeloid leukemia presenting as myeloid sarcoma. Hematol Oncol 30, 34–40 (2012).

13. Begna, K.H., et al. De novo isolated myeloid sarcoma: comparative analysis of survival in 19 consecutive cases. Br J Haematol 195, 413–416 (2021).

14. Bourlon, C., et al. Extramedullary disease at diagnosis of AML does not influence outcome of patients undergoing allogeneic hematopoietic cell transplant in CR1. Eur J Haematol 99, 234–239 (2017).

15. Movassaghian, M., et al. Presentation and outcomes among patients with isolated myeloid sarcoma: a Surveillance, Epidemiology, and End Results database analysis. Leuk Lymphoma 56, 1698–1703 (2015).

16. Fianchi, L., et al. Extramedullary Involvement in Acute Myeloid Leukemia. A Single Center Ten Years’ Experience. Mediterr J Hematol Infect Dis 13, e2021030 (2021).

17. Harris, A.C., et al. Extramedullary relapse of acute myeloid leukemia following allogeneic hematopoietic stem cell transplantation: incidence, risk factors and outcomes. Haematologica 98, 179–184 (2013).

18. Shan, M., et al. Characteristics and transplant outcome of myeloid sarcoma: a single-institute study. Int J Hematol 113, 682–692 (2021).

19. Zorn, K.E., Cunningham, A.M., Meyer, A.E., Carlson, K.S. & Rao, S. Pediatric Myeloid Sarcoma, More than Just a Chloroma: A Review of Clinical Presentations, Significance, and Biology. Cancers (Basel) 15(2023).

20. Wang, C.X., Pusic, I. & Anadkat, M.J. Association of Leukemia Cutis With Survival in Acute Myeloid Leukemia. JAMA Dermatol 155, 826–832 (2019).

21. Engel, N.W., et al. Newly diagnosed isolated myeloid sarcoma-paired NGS panel analysis of extramedullary tumor and bone marrow. Ann Hematol 100, 499–503 (2021).

22. Fouillet, L., et al. A complex mutational profile and a distinct clonal evolution during NPM1 myeloid sarcoma. Leuk Lymphoma 60, 2328–2330 (2019).

23. Greenland, N.Y., et al. Genomic analysis in myeloid sarcoma and comparison with paired acute myeloid leukemia. Hum Pathol 108, 76–83 (2021).

24. Pastoret, C., et al. Detection of clonal heterogeneity and targetable mutations in myeloid sarcoma by high-throughput sequencing. Leuk Lymphoma 58, 1008–1012 (2017).

25. Untaaveesup, S., et al. Genetic alterations in myeloid sarcoma among acute myeloid leukemia patients: insights from 37 cohort studies and a meta-analysis. Front Oncol 14, 1325431 (2024).

26. Zhou, T., et al. Pediatric myeloid sarcoma: a single institution clinicopathologic and molecular analysis. Pediatr Hematol Oncol 37, 76–89 (2020).

27. Clark, W.B., Strickland, S.A., Barrett, A.J. & Savani, B.N. Extramedullary relapses after allogeneic stem cell transplantation for acute myeloid leukemia and myelodysplastic syndrome. Haematologica 95, 860–863 (2010).

28. Yoshihara, S., Ando, T. & Ogawa, H. Extramedullary relapse of acute myeloid leukemia after allogeneic hematopoietic stem cell transplantation: an easily overlooked but significant pattern of relapse. Biol Blood Marrow Transplant 18, 1800–1807 (2012).

29. Christopher, M.J., et al. Immune Escape of Relapsed AML Cells after Allogeneic Transplantation. N Engl J Med 379, 2330–2341 (2018).

30. Tohalori, C., et al. Immune signature drives leukemia escape and relapse after hematopoietic cell transplantation. Nat Med 25, 603–611 (2019).

31. Pagliuca, S., et al. Leukemia relapse via genetic immune escape after allogeneic hematopoietic cell transplantation. Nat Commun 14, 3153 (2023).

32. Stolzel, F., et al. Clonal evolution including partial loss of human leukocyte antigen genes favoring extramedullary acute myeloid leukemia relapse after matched related allogeneic hematopoietic stem cell transplantation. Transplantation 93, 744–749 (2012).

33. Burd, A., et al. Precision medicine treatment in acute myeloid leukemia using prospective genomic profiling: feasibility and preliminary ehicacy of the Beat AML Master Trial. Nat Med 26, 1852–1858 (2020).

34. Genomes Project, C., et al. A global reference for human genetic variation. Nature 526, 68–74 (2015).

35. Sango, J., et al. RAS-mutant leukaemia stem cells drive clinical resistance to venetoclax. Nature 636, 241–250 (2024).

36. Bruserud, O., Selheim, F., Hernandez-Valladares, M. & Reikvam, H. Monocytic Diherentiation in Acute Myeloid Leukemia Cells: Diagnostic Criteria, Biological Heterogeneity, Mitochondrial Metabolism, Resistance to and Induction by Targeted Therapies. Int J Mol Sci 25(2024).

37. Bennett, J.M., et al. Proposed revised criteria for the classification of acute myeloid leukemia. A report of the French-American-British Cooperative Group. Ann Intern Med 103, 620–625 (1985).

38. Miller, C.A., et al. Visualizing tumor evolution with the fishplot package for R. BMC Genomics 17, 880 (2016).

39. Islam, S.M.A., et al. Uncovering novel mutational signatures by de novo extraction with SigProfilerExtractor. Cell Genom 2, None (2022).

40. Sondka, Z., et al. The COSMIC Cancer Gene Census: describing genetic dysfunction across all human cancers. Nat Rev Cancer 18, 696–705 (2018).

41. Bamford, S., et al. The COSMIC (Catalogue of Somatic Mutations in Cancer) database and website. Br J Cancer 91, 355–358 (2004).

42. Alexandrov, L.B., et al. Clock-like mutational processes in human somatic cells. Nature genetics 47, 1402–1407 (2015).

43. Crucitti, L., et al. Incidence, risk factors and clinical outcome of leukemia relapses with loss of the mismatched HLA after partially incompatible hematopoietic stem cell transplantation. Leukemia 29, 1143–1152 (2015).

44. Vago, L., et al. Loss of mismatched HLA in leukemia after stem-cell transplantation. N Engl J Med 361, 478–488 (2009).

45. Wu, H., et al. Assessment of Patient-Specific Human Leukocyte Antigen Genomic Loss at Relapse After Antithymocyte Globulin-Based T-Cell-Replete Haploidentical Hematopoietic Stem Cell Transplant. JAMA Netw Open 5, e226114 (2022).

46. Biassoni, R., et al. Amino acid substitutions can influence the natural killer (NK)-mediated recognition of HLA-C molecules. Role of serine-77 and lysine-80 in the target cell protection from lysis mediated by “group 2” or “group 1” NK clones. J Exp Med 182, 605–609 (1995).

47. Sobecks, R.M., et al. Influence of killer immunoglobulin-like receptor/HLA ligand matching on achievement of T-cell complete donor chimerism in related donor nonmyeloablative allogeneic hematopoietic stem cell transplantation. Bone Marrow Transplant 41, 709–714 (2008).

48. Stringaris, K., et al. Donor KIR Genes 2DL5A, 2DS1 and 3DS1 are associated with a reduced rate of leukemia relapse after HLA-identical sibling stem cell transplantation for acute myeloid leukemia but not other hematologic malignancies. Biol Blood Marrow Transplant 16, 1257–1264 (2010).

49. Venstrom, J.M., et al. HLA-C-dependent prevention of leukemia relapse by donor activating KIR2DS1. N Engl J Med 367, 805–816 (2012).

50. Dufva, O., et al. Immunogenomic Landscape of Hematological Malignancies. Cancer Cell 38, 380–399 e313 (2020).

51. Chauhan, P.S., et al. Mutation of NPM1 and FLT3 genes in acute myeloid leukemia and their association with clinical and immunophenotypic features. Dis Markers 35, 581–588 (2013).

52. Cauchy, P., et al. Chronic FLT3-ITD Signaling in Acute Myeloid Leukemia Is Connected to a Specific Chromatin Signature. Cell Rep 12, 821–836 (2015).

53. Ferrito, N., et al. Biomarker Landscape in RASopathies. Int J Mol Sci 25(2024).

54. Loh, M.L. Recent advances in the pathogenesis and treatment of juvenile myelomonocytic leukaemia. Br J Haematol 152, 677–687 (2011).

55. Quek, C.W.N., et al. Leukemia cutis in a child with juvenile myelomonocytic leukemia presenting with Sweet syndrome-like lesions and a history of multiple juvenile xanthogranulomas. JAAD Case Rep 6, 1138–1140 (2020).

56. Pylayeva-Gupta, Y., Grabocka, E. & Bar-Sagi, D. RAS oncogenes: weaving a tumorigenic web. Nat Rev Cancer 11, 761–774 (2011).

57. Yaeger, R., et al. RAS mutations ahect pattern of metastatic spread and increase propensity for brain metastasis in colorectal cancer. Cancer 121, 1195–1203 (2015).

58. de Braud, F., et al. Initial Evidence for the Ehicacy of Naporafenib in Combination With Trametinib in NRAS-Mutant Melanoma: Results From the Expansion Arm of a Phase Ib, Open-Label Study. J Clin Oncol 41, 2651–2660 (2023).

59. Cox, A.D., Der, C.J. & Philips, M.R. Targeting RAS Membrane Association: Back to the Future for Anti-RAS Drug Discovery? Clin Cancer Res 21, 1819–1827 (2015).

60. Punekar, S.R., Velcheti, V., Neel, B.G. & Wong, K.K. The current state of the art and future trends in RAS-targeted cancer therapies. Nat Rev Clin Oncol 19, 637–655 (2022).

61. Shetu, S.A. & Bandyopadhyay, D. Small-Molecule RAS Inhibitors as Anticancer Agents: Discovery, Development, and Mechanistic Studies. Int J Mol Sci 23(2022).

62. Song, Y., et al. Targeting RAS-RAF-MEK-ERK signaling pathway in human cancer: Current status in clinical trials. Genes Dis 10, 76–88 (2023).

63. Ram, T., et al. MEK inhibitors in cancer treatment: structural insights, regulation, recent advances and future perspectives. RSC Med Chem 14, 1837–1857 (2023).

64. Pagliuca, S., et al. Comprehensive Transcriptomic Analysis of VISTA in Acute Myeloid Leukemia: Insights into Its Prognostic Value. Int J Mol Sci 23(2022).

65. Robinson, J.T., et al. Integrative genomics viewer. Nat Biotechnol 29, 24–26 (2011).

66. Gurnari, C., et al. Clinical and Molecular Determinants of Clonal Evolution in Aplastic Anemia and Paroxysmal Nocturnal Hemoglobinuria. J Clin Oncol 41, 132–142 (2023).

67. Pagliuca, S., et al. Molecular landscape of immune pressure and escape in aplastic anemia. Leukemia 37, 202–211 (2023).

