## Supplementary Appendix and Figures for "Genomic and Immunogenomic Profiling of Extramedullary Acute Myeloid Leukemia Reveals Actionable Clonal Branching and Frequent Immune Editing"

**Running title:** Clonal branching and Immunoediting in eAML

**Authors:** Clément Collignon[1,2], Tucker Hansen[1], Colin Hercus[3], Marianna B. Ruzinova[4], Gabrielle Roth Guepin[2], Caroline Bonmati[2], Marie Thérèse Rubio[2,5], Pierre Feugier[2], Mélanie Gaudfrin[5], Hervé Sartelet[6] Marion Divoux[7], Marc Muller[7], Sharon Heath[1], Geoffrey Uy[1], David Spencer[1], David Chen[8], Simona Pagliuca*[2,5] & Francesca Ferraro*[1].

**Affiliations:**
1 Division of Oncology – Hematologic Malignancies, Transplant and Cellular Therapy, Washington University in Saint Louis
2 Hematology Department, Nancy University Hospital, Vandoeuvre-lès-Nancy, France
3 Novocraft Technologies Sdn Bhd, Kuala Lumpur, Malaysia.
4. Department of Pathology and Immunology, Washington University in Saint Louis
5 UMR 7365, IMoPA, University of Lorraine, Vandoeuvre-lès-Nancy, France
6 Department of Pathology, Nancy University Hospital, Vandoeuvre-lès-Nancy, France
7 Genetic Department, Nancy University Hospital, Vandoeuvre-lès-Nancy, France
8 Division of Dermatology, Washington University in Saint Louis

**Patients' clinical summaries**

**204235 (EMD 1)**

2022 ICC Classification: **AML with t(8;16)(p11.2;p13.3)/*KAT6A::CREBBP*, therapy-related**

2022 ELN risk classification: **Adverse**

40-year-old female with history of triple negative breast cancer, stage IIIc, treated with neoadjuvant chemotherapy on the Keynote 522 (weekly carboplatin and paclitaxel x 4 cycles + and adriamycin/cyclophosphamide every 3 weeks for 4 cycles + pembrolizumab), followed by right radical mastectomy. Two months after completing the treatment, she developed a skin rash and was diagnosed with *myeloid sarcoma*. Concomitant bone marrow biopsy showed multilineage dysplasia but no excess blasts. Karyotype 46,XX,t(8;16)(p11.2;p13.3). FISH showing *MYST3/CREBBP* at 3.5% and molecular showing *PPM1D* frameshift at 8%. She received one cycle of Vyxeos induction. A PET scan at count recovery showed multiple new lung nodules and mediastinal lymphadenopathies. Biopsy of the lesions showed metastatic breast cancer. She was started on capecitabine with progressive disease, followed by eribulin with further progression of the metastatic breast cancer. She transitioned to hospice care and expired 5 months after the diagnosis of myeloid sarcoma and one year and 4 months after the diagnosis of breast cancer.

**208468 (EMD 2)**

2022 ICC Classification: **AML with mutated TP53, progressing from MDS**

2022 ELN risk classification: **Adverse**

65-year-old female diagnosed with secondary AML from high risk MDS EB-1, previously treated with 3 cycles of azacytidine and magrolimab. At AML progression blood counts showing WBC 0.6 with 40% circulating blasts at flow cytometry, Hb of 8.1 and platelet of 22,000. Bone marrow hypercellular with 50% blasts. Karyotype was complex 43~48,XX,?inv(3)(p21p23),del(5)(q23q31),-6,-7,der(8)t(5;8)(p12;p11.1),+16,+1~4mar[cp20]. FISH with 5q-, 7q-, +22q, -13q, and monosomy 13. Molecular showing *IDH1* at 2%, *PTPN11* at 34%, *TP53* at 97%, *SRSF2* at 2%. She received Vyxeos, with a day 30 bone marrow showing persistent disease. She was started on 10 days decitabine and venetoclax with progressive disease and appearance of *leukemia cutis*. She was admitted and received DART on clinical trial. Her overall condition continued to worsen while inpatient and she transitioned to hospice. She expired 5 months after progression to AML for complications of refractory leukemia.

**290344 (EMD 4)**

2022 ICC Classification: **AML not otherwise specified**

2022 ELN risk classification: **Intermediate**

62-year-old female presenting with an epidural mass consistent with myeloid sarcoma at tissue biopsy. Bone marrow concomitant with no excess blasts, normal karyotype and FISH studies. Molecular on bone marrow was not performed. Molecular on myeloid sarcoma showed *DNMT3A*, *NRAS*, *ARID2*, *BCORL1* and MSH6 missense mutations. Patient had a past medical history of acute myeloid leukemia, onset 13 years prior to current presentation, with trisomy 11 in bone marrow at diagnosis. She received sib-allo 12 years prior to myeloid sarcoma presentation. Counts at myeloid sarcoma diagnosis were: WBC 18.5 with no circulating blasts, Hb 11.7 and platelets 433,000. Myeloid sarcoma STR showing recipients cells, consistent with relapsed disease. She received one cycle of 7+3 with PET/CT showing remission of epidural masses. She continued care at an outside hospital, due to insurance issue. She received a cycle of intermediate dose cytarabine consolidation followed by a second allogeneic transplant MUD and expired one year and five months after recurrence of her AML. Circumstances of death are not available due to out of network care.

**297155 (EMD5)**
2022 ICC Classification: **AML with mutated NPM1**
2022 ELN risk classification: **Favorable**
41-year-old male diagnosed with AML (FAB M5). WBC at diagnosis 4.7, with 7% circulating blasts. Hb 9.4 and platelet 377,000. Bone marrow biopsy with 90% cellularity and 21% blasts/ Karyotype and FISH normal. Molecular showing *DNMT3A* R882H 28%, *NPM1* W288Cfs*12 19% and *FLT3-TKD* 4%. Risk category: favorable. He underwent 7+3 induction, day 30 bone marrow was MRD negative by molecular testing. He received 4 cycles of HiDAC consolidation with midostaurin with no initial plan for transplant in CR1. He relapsed with *myeloid sarcoma* of the left testicle, which was treated with orchiectomy and radiation. He was admitted to undergo further treatment with salvage chemotherapy, due to development of leukocytosis and multiple skin nodules consistent with *leukemia cutis*. Upon admission he developed altered mental status and a CT scan showed intraparenchymal and intraventricular brain hemorrhage. His neurological status continued to deteriorate rapidly, and he expired 8 months after the initial AML diagnosis.

**319693 (EMD 6)**
2022 ICC Classification: **AML with myelodysplasia-related gene mutations**
2022 ELN risk classification: **Adverse**
80-year-old male with a diagnosis of AML (FAB M2) and *leukemia cutis* at diagnosis. Bone marrow at diagnosis showed 38% blasts. Blood count at diagnosis: WBC 17,500 with 68% circulating blasts, Hb 4.8, plt 29,000. Karyotype 47,XY,+8[20]. FISH with trisomy *RUNX1T1* (8q) - 94%. Molecular with *IDH2* at 45%, *SRSF2* at 35% and *STAG2* at 88%. Risk category: adverse. He received low dose Vyxeos and venetoclax. Bone marrow biopsy at count recovery with morphologic remission, FISH with persistent trisomy *RUNX1T1* (8q) at 66%. Patient transitioned to palliative care at an outside hospital. He expired 4 months after his initial leukemia diagnosis.

**375074 (EMD 7)**
2022 ICC Diagnosis: **AML with mutated NPM1 and myelodysplasia-related cytogenetic abnormalities**
2022 ELN risk classification: **Adverse**
69-year-old diagnosed with AML (FAB M5). Blood counts at diagnosis with WBC 67,000 with 63% circulating blasts, Hb 8.9 Platelet 37,000. Bone marrow with >90% cellularity and 78% monocytic blasts. Cytogenetic showing 46,XY, del(1)(p36.2), -7. Molecular testing with *NPM1* W288Cfs*12 30%, *IDH2* R140Q 48%, *SRSF2* P59R 44%, *TET2* V1180D 77% mutations.
He received induction with 7 + 3 + uprolesan on clinical trial. He then underwent consolidation with intermediate dose cytarabine and uprolesan for three cycles achieving complete remission. He relapsed 3 years and 5 months post diagnosis. He received one cycle of azacytidine and venetoclax complicated by pancytopenia and development of *leukemia cutis.* He was admitted to the hospital for high dose cytarabine but develop altered mental status and abdominal distention. He was transitioned to hospice care and died 1334 days after diagnosis for complication of relapsed disease and sepsis.

**442721 (EMD 8)**
2022 ICC Classification: **AML with myelodysplasia-related cytogenetic abnormality**

2022 ELN risk classification: **Adverse**
64-year-old male diagnosed with AML with myelodysplastic related changes. (AML w/MRC). Blood counts at diagnosis showing WBC 2.7 with 23% circulating blasts, Hb 7.9 and platelets 225,000. Bone marrow cellularity 60% with 30% blasts. Cytogenetic with complex karyotype 46,XY,der(3)?del(3)(q12q13)ins(3;?)(q11.2;?), -5,del(5)(q13q33),del(5)(q31q35),del(7)(q22q34),+mar[cp14]/46,XY[6]. FISH with monosomy 5 and del5q. Molecular diagnostic with *ASXL1*, *BCOR*, *RUNX1*, *NF1*. He received Vyxeos induction. Day +14 marrow showing persistent disease. He received a second Vyxeos reinduction with persistent disease at count recovery. He underwent salvage with FLAG-IDA and achieved morphologic remission. He underwent haploidentical transplant and relapsed 2 years and seven months after transplant with *myeloid sarcoma* without bone marrow involvement. He received radiation to the myeloid sarcoma and underwent a second allogeneic MMUD (9/10). His day 100 bone marrow was MRD positive. He was started on 5 days azacytidine with plan for a DLI. However, he was hospitalized in the setting of pancytopenia and dyspnea. He was diagnosed with fungal pneumonia. His clinical status continued to deteriorate, and he was transitioned to hospice. He expired 4 years and 5 months after his initial AML diagnosis.

**464271(EMD 9)**
2022 ICC Classification: **AML KMT2A-rearranged KMT2A::ELL**
2022 ELN risk classification: **Adverse**
47-year-old female with diagnosis of AML (FAB M5). Blood count at diagnosis showing WBC 2,300 with 20% circulating blasts. Hb 7.2 plt 100,000. Bone marrow biopsy with increased cellularity and 94% blasts. Karyotype with 46,XX,t(11;19)(q23;p13.3)[20]. Molecular with *ASXL1* G646Wfs*12 13%. Risk category: adverse. She received induction chemotherapy with 7+3. Day 30 bone marrow showed complete remission. She received one cycle of HiDAC consolidation, followed by allogeneic sibling transplant. 3 months after transplant, her bone marrow FISH was positive at 2.5%. She received 2 cycles of decitabine and venetoclax with MRD negative remission. This was followed by a DLI and one cycle of 5 days decitabine, after which she developed progressive disease with 15% blasts in the bone marrow, *myeloid sarcoma* (psoas muscle) and central nervous system involvement. She received FLAG-IDA salvage and LP with MTX and AraC, with CNS clearance and PET negativity. She was subsequently admitted for a second allogeneic transplant, which was complicated by septic shock and diffuse alveolar hemorrhage. She expired 21 days after her second transplant and one year after initial AML diagnosis.

**469908 (EMD 10)**
2022 ICC Diagnosis: **AML with mutated NPM1**
2022 ELN risk classification: **Favorable**
76-year-old male diagnosed with AML (FAB M5). He presented with pancytopenia and *leukemia cutis*. Blood counts at diagnosis: WBC 2,100, Hb 7.7 and platelet 120,000. Bone marrow biopsy at diagnosis with hypercellular marrow and 80% blasts. Karyotype normal, FISH negative. Molecular showing NPM1 W288Cfs*12 15%, *DNMT3a* c.1667+1G>T 19% and *SF3B1* K666N 7%. Risk category: favorable. He was started on treatment on the BLAST trial (azacytidine, venetoclax and pembrolizumab). Bone marrow biopsy after one cycle with CR, MRD negative. He underwent 6 more cycles of treatment, without pembrolizumab after cycle 5. After cycle 3 bone marrow showed morphological clearance but MRD pos for DNMT3A and SF3B1 mutations. His BM after cycle 7 showed relapsed disease with 18% blasts. He was admitted for intermediate dose cytarabine. The treatment was complicated by septic shock and death 8 months after initial diagnosis of leukemia.

**562600 (EMD 12)**
2022 ICC Classification: **AML, with mutated NPM1**
2022 ELN risk classification: **intermediate**
60-year-old male diagnosed with AML with myelodysplastic related changes (AML w/MRC). Blood counts at diagnosis with WBC 30,400 with 31% circulating blasts, Hb 7.2, platelet 50,000. Bone marrow with dysplasia and 20% blasts. Cytogenetic with normal karyotype. Molecular showing *NRAS* G12D at 5%, *DNMT3A* R882G at 44%, *FLT3 -ITD* exon 14 insertion (VAF not available), *CEBPA* H24Afs*12 12% and Q41* 33%, *RAD21* G237C at 4%, *IDH2* R140W at 13%, *NPM1* W288Cfs*12 41%. He received induction chemotherapy with 7 + 3 and one cycle of HiDAC maintenance. He tolerated HiDAC poorly with several ICU admissions. He was placed on gilteritinib maintenance. He relapsed one year after diagnosis with *leukemia cutis* and bone marrow involvement. He received one cycle of Decitabine + Venetoclax and subsequently enrolled in a clinical trial with SEL120 and was in remission for 6 months. He relapsed again with systemic disease and was to start on Syndex trial, but his clinical status deteriorated with development of hypoxic respiratory failure. He transitioned to hospice and died 2.5 years after initial diagnosis.

**585305 (EMD 13)**
2022 ICC Classification: **AML with myelodysplasia-related gene mutations**
2022 ELN risk classification: **Adverse**
43-year-old male with AML and *leukemia cutis* at diagnosis. Blood counts at presentation WBC 25,500 with 63% circulating blasts, Hb 7.9 platelet 26,000. BM at diagnosis showed AML with monocytic differentiation (FAB M5), with 41% blasts. Cytogenetic showed normal karyotype and molecular showed *FLT3-ITD* at 9% VAF and *RUNX1* pA338Rfs*262 at 32%. Risk category: adverse. He was started on induction chemotherapy with 7+3 and midostaurin. Day 14 bone marrow showed persistent disease. Lumbar puncture was negative for central nervous system involvement. He received reinduction with azacytidine, venetoclax and gilteritinib and achieved remission after 3 cycles. He was unable to proceed to transplant due to poor social situation and recurrent infections. He was started on azacytidine maintenance orally 300 mg PO day1-14, which he did not tolerate. Oral maintenance was changed to gilteritinib, on which he developed progressive pancytopenia and relapsed disease. On 06/27/2024 he is 752 days post diagnosis alive receiving palliative care.

**615755 (EMD 15)**
2022 ICC Classification: **AML with *NUP98* and other partners and mutated NPM1**
2022 ELN risk classification: **Favorable**
60-year-old male with diagnosis of AML (FAB M5) and *leukemia cutis* at presentation. Blood counts at diagnosis with WBC 125,000 with 11% circulating blasts, Hb 7.9, platelet 50,000. BM at diagnosis with 90% cellularity and 40% blasts. Cytogenetic showing 46,XY, t(11;17)(p15;q12)[20] with atypical NUP98 rearrangement in 90.5% of cells by FISH. Molecular studies showing NPM1 *W288Cfs*12* 37%, *PTPN11* G60V 9% and S502L at 20%, *TET2* M1388Afs*12 39% and E1923* 42%, *NF1* S1838C 47%. He received induction with 7 + 3. Day 14 bone marrow chemoablated. Day 30 BM with morphologic remission, but molecular MRD positive. Underwent 10/10 MUD stem cell transplant and he is currently 3 years and 1-month post-transplant, in complete remission and off immunosuppression.

**664428 (EMD 17)**
2022 ICC Classification: **AML with mutated NPM1, progressing from MDS**
2022 ELN risk classification: **Favorable**

67-year-old male diagnosed with secondary AML (from MDS, diagnosed 5 months prior) and leukemia cutis at AML progression (FAB M4). At AML progression, blood counts showed: WBC 56,600 with 74% circulating blasts, Hb 7.0, plt 22,000. Bone marrow biopsy with hypercellular marrow and 53% blasts. Karyotype 46,XY,der(10)t(3;10)(q21;p11.2)[10]/46,XY[10]. FISH with trisomy of MECOM (3q) 34.5%. Molecular studies showing NRAS G12D at 6%, TET2 C1211Y at 46% and F1309L 45%, NPM1 W288Cfs*12 at 16%, SRSF2 P95R at 49% , RAD21 intron splice variant at 36% and KRAS G12R at 42. Upon AML progression he received Vyxeos. His BM at count recovery showed morphological remission with MRD 0.3%. He received 5 days decitabine for 3 cycles, complicated by pancytopenia and recurrent GI bleed, progressive disease, with worsening leukemia cutis. He transitioned to hospice and expired 7 months after progression to AML.

**704854 (EMD 18)**
2022 ICC Classification: **AML with myelodysplasia-related gene mutations**
2022 ELN risk classification: **Adverse**
55 years old male with diagnosis of AML (FAB M4). Blood count at diagnosis with WBC 3,900 and no circulating blasts, Hb 8.3 and platelets 19,000. Bone marrow biopsy with hypercellular marrow and 32% blasts. Karyotype failed due to insufficient metaphases, FISH negative. Molecular with *DNMT3A* R889C 42%, DNMT3A *R598** 45%, GATA2 *T387_K389del* 30%, EZH2 *I744T* 5%, *ASXL1* A640Gfs*60 and P808Lfs*10 at 4 and 41% VAF respectively, and *PTPN11* G503V 25%. Risk category: adverse. He received 7+3 induction. Day 14 BM chemoablated. Day 30 bone marrow with morphologic remission but positive MRD. He relapsed after 1 cycle of HIDAC consolidation and was started on 10 days Decitabine and Venetoclax with morphologic response, after which he received allogeneic transplant on IOMAB study. He relapsed early post-transplant and received DLI followed by 5 days Decitabine and Venetoclax. He achieved complete remission with 100 percent donor engraftment. He relapsed again 334 days post-transplant with isolated *leukemia cutis* to his axilla without evidence of systemic relapse. He received radiation to the axilla but rapidly developed additional myeloid sarcoma localizations and expired shortly thereafter for complication of recurrent leukemia.

**775682 (EMD 19)**
2022 ICC Classification: **AML with mutated NPM1**
2022 ELN risk classification: **Favorable**
83-year-old male diagnosed with AML (FAB M5). Blood counts at diagnosis with WBC 119,000 and 67% circulating blasts, Hb 10 and platelet 46,000. Bone marrow cellularity 80% with 91% blasts. Karyotype was normal. Molecular showed *NPM1* W288Cfs*12 44%, *CBL* p.C404Y 70% and *IDH2* p.R140Q 51%. Risk category: favorable. He was hospitalized and received one cycle of azacytidine and venetoclax and was then transitioned to enasidenib. He remained pancytopenic throughout, and developed *myeloid sarcoma* on enasidenib 1.5 months after diagnosis. He was readmitted with altered mental status, MRSA bacteremia, acute kidney injury and pneumonia and died of complications leukemia and septic shock 3 months post diagnosis.

**816461 (EMD 21)**
2022 ICC Classification: **AML with other *KMT2A* rearrangement**
2022 ELN risk classification: **Adverse**
60-year-old female with history of marginal zone lymphoma transformed to DLBCL status post treatment with R-CHOPx6, BRx2, RICEx1 and MUD transplant, followed by 3 DLI due

to relapsed DLBCL, who developed *leukemia cutis* with concomitant bone marrow involvement by donor-derived AML (FAB M5) with 56% blasts. Blood count at leukemia diagnosis WBC 3.9 with 24% circulating blasts, Hb 10.9 and platelet 102,000. Karyotype showing /46,XY,t(11;19)(q23.3;p13.3)[14]/46,XY[6], FISH with 100% donor cells and KMT2A rearrangement - 33.5%. She received 2 cycles of decitabine and venetoclax with no response, and bone marrow blasts increase to 70% and persistent *leukemia cutis*. She was transitioned to hospice care. She expired 6 months after the AML diagnosis.

**895870 (EMD 22)**
2022 ICC Classification: **AML with mutated *NPM1***
2022 ELN risk classification: **Favorable**
63-year-old female diagnosed with AML, FAB M2. Blood counts at presentation were WBC 39.8, with 89% circulating blasts, Hb 8.9 and platelets 25,000. A bone marrow aspiration demonstrated 90 % cellularity with 62% blasts. Cytogenetic showed normal karyotype 46,XX[20]. Molecular diagnostic studies showed NPMc W288Cfs*12 28%, *IDH2* R140Q 48%, *ASXL1* G646Wfs*12 11%, *FLT3-TKD* V592G 27%, *SRSF2* R94dup 45%, *STAG2* A956Ffs*4 9%. She received standard of care induction chemotherapy with 7+3. A day 14 bone marrow was chemoablated. A day 28 bone marrow showed complete remission. She received consolidation with 3 cycles of intermediate dose cytarabine 2 grams/m2. End of treatment biopsy showed complete remission and MRD negative by flow. She relapsed with leukocytosis and *leukemia cutis* 6 months after completing treatment, bone marrow biopsy at relapse showed 80% cellularity with 55% blasts. She received enasidenib and azacytidine, achieved morphologic remission after cycle 1 but had progressive disease after cycle 2. She was transitioned to azacytidine and venetoclax, but developed cellulitis and had a fall with head trauma. Her performance status deteriorated and she elected to transition to Hospice. She expired 18 months after initial diagnosis.

**Nancy_2 (EMD 23)**
2022 ICC Classification: **AML with t(8;21)(q22;q22.1)/*RUNX1::RUNX1T1***
2022 ELN risk classification: **Favorable**
57-year-old male diagnosed with AML, FAB M1. Blood counts at presentation were WBC 45.84 hemoglobin 9.5, platelets 59, with 88% circulating blasts. A bone marrow aspiration demonstrated 93% blasts. Cytogenetics showed a 45,X,-Y, t(8;21)(q22;q22) karyotype. Molecular diagnostic studies included AML1::ETO transcript positive, KIT N822K 37% mutation. Risk category: favorable. Initial therapy consisted of daunorubicin and cytarabine induction. A bone marrow aspiration at midcycle demonstrated an ablated marrow. A repeat marrow at count recovery demonstrated remission. Post-remission chemotherapy consisted of 3 cycles of high dose cytarabine consolidation. Isolated extramedullary relapse was documented 14 months post diagnosis in biopsied clinically apparent isolated *extramedullary paravertebral C8-T1 lesion*. Medullary disease reevaluation showed a normal appearing bone marrow with AML1::ETO detected transcript and extramedullary investigation with molecular studies with FLT3 V581_K602dup 34.4% mutation. Treatment consisted with mitoxantrone, cytarabine, gemtuzumab ozagamycin salvage resulting in CR2 complemented with gilteritinib maintenance. He underwent haploidentical stem cell transplant followed by gilteritinib maintenance. He remains in second complete remission 17 months after relapse.

**Nancy_3 (EMD 24)**
2022 ICC Classification: **AML, not otherwise specified**
2022 ELN risk classification: **Intermediate**

2-year-old male diagnosed with bony palate *myeloid sarcoma*. Blood counts at presentation were WBC 9.85 hemoglobin 10.1, platelets 573. Bone marrow aspiration with cytogenetics, and molecular diagnostic studies were normal. NGS performed showed IKZF1 R184W 3% on bone marrow sample and IKZF1 R184W 41% and ETV6 R105L 40% mutations on extramedullary lesion, morphology showed a monocytic differentiation and immunophenotyping was CD33+, CD34+, CD117+, CD123dim, MPO+/-, CD19+/-, CD22-. Risk category: intermediate. Initial therapy consisted of mitoxantrone, cytarabine, gemtuzumab ozogamycine induction with intrathecal cytarabine injection per MyeChild protocol. A repeat palate biopsy at count recovery demonstrated persistent disease with 4% myeloid blasts on bone marrow aspiration. Treatment consisted of idarubicine, fludarabine and high dose cytarabine salvage resulting in first complete remission. Post-remission chemotherapy consisted of 1 cycle of fludarabine and HiDAC consolidation. He underwent matched unrelated donor allogenic stem cell transplant in CR1. Isolated extramedullary relapse was documented 9 months post diagnosis in biopsied clinically apparent isolated extramedullary bony palate lesion. Marrow aspiration showed a normal appearing bone marrow and extramedullary investigation with NGS revealed IKZF1 R184W 11%. Treatment consisted with HiDAC and venetoclax salvage associated with 18Gy proton-therapy resulting in CR2. Post-remission treatment consisted of 1 cycle of azacytidine and venetoclax, and 1 cycle of azacytidine consolidation. Medullary relapse was documented 15 months post diagnosis. Treatment consisted with 1 cycle of cytarabine and venetoclax associated with 1 DLI as palliative care. He expired from progressive disease 17 months post diagnosis.

**Nancy_13 (EMD 27)**
<u>2022 ICC Classification:</u> **AML, not otherwise specified**
<u>2022 ELN risk classification</u>: **Intermediate**
56-year-old female diagnosed with AML M5. Blood counts at diagnosis with WBC 8,600 with 52% circulating blasts, Hb 9.9 Platelet 183,000. Bone marrow with 84% blasts. Cytogenetic with normal karyotype and molecular diagnostic with no abnormalities. Risk category: intermediate. She received induction chemotherapy with daunorubicin and cytarabine. Bone marrow aspiration at count recovery showed no response with 65% blasts. Salvage therapy consisted of 1 cycle of HiDAC, with refractory disease at count recovery evaluation and emergence of lesions suspected of *leukemia cutis*. She received 1 cycle of azacytidine + venetoclax with no response. Further salvage therapy consisted of mitoxantrone + cytarabine + gemtuzumab ozogamycin. She achieved CR1 and underwent matched related transplant 185 days after diagnosis. She relapsed with extramedullary involvement of skin, breast, kidney, peritoneum, retro-peritoneum, and pancreas suspected on TDM and expired 498 days after AML diagnosis of acute cholangitis and pancreatitis.

**Nancy_14 (EMD 28)**
<u>2022 ICC Classification:</u> **AML with myelodysplasia-related gene mutations**
<u>2022 ELN risk classification</u>: **Adverse**
71-year-old male diagnosed with AML M2. Blood counts at presentation were WBC 24.27 hemoglobin 9.6, platelets 150, with 70% circulating blasts. A bone marrow aspiration demonstrated 54% blasts. Cytogenetics showed a 47,XY,+13 karyotype. Molecular diagnostic highlighted ASXL1 G646Wfs*12 38.8%, CEBPA K171* 50.9%, SRSF2 P95H 48.3%, TET2 E1323Ffs*13 46.0% and TET2 P1194L 48.8% mutations. Risk category: adverse. Work-up also revealed an *asymptomatic extramedullary involvement of ileo-caecal valve*. Initial therapy consisted of azacytidine and venetoclax induction. A repeat marrow at the end of the

first course showed a partial response with 15% blasts. Azacytidine and venetoclax were stopped after 2 more courses. He expired from progressive disease 7 months post diagnosis.

**Nancy_15 (EMD 29)**
2022 ICC Classification: **AML with myelodysplasia-related cytogenetic abnormalities**
2022 ELN risk classification: **Adverse**
31-year-old female with diagnosis of AML associated with blastic pericarditis and pleurisy. Blood counts at diagnosis with WBC 16,000 with 45% circulating blasts, Hb 11.2 Platelet 160,000. Bone marrow blasts percentage is not available. Cytogenetic showing complex karyotype. Molecular testing is not available. Risk category: adverse. Induction therapy consisted of idarubicine and cytarabine. Day 30 bone marrow showed CR1. She received 1 cycle of HiDAC consolidation with relapsed disease at count recovery associated with *blastic meningitidis* suspicion. Disease was refractory against cytoreduction agents. She expired 85 days post diagnosis after CRA post-pericardic drain.

**Nancy_16 (EMD 30)**
2022 ICC Classification: **AML with inv(16)(p13.1q22) or t(16;16)(p13.1;q22)/*CBFB::MYH11***
2022 ELN risk classification: **Favorable**
66-year-old male diagnosed with AML M5a associated with suspected leukemia cutis. Blood counts at diagnosis with WBC 353,000 with 82% circulating blasts, Hb 3.9 Platelet 36,000. Bone marrow with 88% blasts. Cytogenetic showing 46,XY, inv(16)(p13q22) confirmed with FISH. Molecular testing with KIT D816V 45% mutation. Risk category: favorable. Initial therapy consisted of idarubicine, cytarabine, lomustine. Day 30 bone marrow showed complete remission. He received 3 cycles of IDAC consolidation with allogenic sibling transplant in CR1. 3 months after transplant, molecular relapse was treated with 1 cycle of azacytidine and midostorin and 3 cycles of azacytidine. 17 months post diagnosis, cerebellar syndrome led to perform MRI pointing *one cerebellar and multiple cerebral masses consistent with AM*L at tissue biopsy with FLT3-ITD mutation, associated with marrow molecular relapse. He received radiotherapy + venetoclax + gilteritinib. He expired of progressive disease 584 days post-transplant and 24 months post-diagnosis.

**Nancy_17 (EMD 31)**
2022 ICC Classification: **AML with mutated *NPM1* and myelodysplasia-related gene mutations**
2022 ELN risk classification: **Favorable**
71-year-old male diagnosed with AML associated with documented leukemia cutis (biopsied, not available for sequencing). Blood counts at diagnosis with WBC 4,300 with 3% circulating blasts, Hb 8.2 Platelet 54,000. Bone marrow with 83% blasts. Cytogenetic showing 46,XY,del(20)(q11;q13)[18]/47,XY,+8 [8]/46,XY[5]. Molecular showing IDH2 R140Q 34.9%, SRSF2 P95H 32%, NRAS Q61L 2.1%, NPM1 W288Cfs*12 3.2%, RUNX1 R320*1.3%, TET2 H1912R 3.3%, DNMT3A F354V 2.9%, EZH2 D189del 3.5%. Risk category: intermediate. Initial therapy consisted of idarubicine, cytarabine, lomustine. Day 30 bone marrow showed complete remission. Without compatible donor for allogenic transplant in CR1, he received 2 reinduction with idarubicine, cytarabine followed by 4 cycles of oral azacytidine. Testicular hypertrophy led to document *myeloid sarcoma extramedullary relapse* on orchidectomy histology, associated with medullary cytogenetic relapse 823 days post-diagnosis. He received 1 cycle of azacytidine + venetoclax resulting in CR2 followed with 4 cycles of azacytidine + venetoclax. He expired of progressive disease 1065 days post diagnosis.

**Nancy_18 (EMD32)**
2022 ICC Classification: **AML with mutated NPM1**
2022 ELN risk classification: **Intermediate**
59-year-old male diagnosed with AML M4. Blood counts at diagnosis with WBC 66,000 with 21% circulating blasts, Hb 9 Platelet 75,000. Bone marrow with 23% blasts. Karyotype and FISH normal. Molecular showing NPM1 type A 35.5%, FLT3-ITD Y599_E604dup 19.8%, FLT3-ITD Y597_P606dup 3.2%, DNMT3A Q692Tfs*21 VAF 43.8 %. Risk category: Intermediate. Initial therapy consisted of 2 cycles of daunorubicine, cytarabine, gilteritinib on HOVON 156 clinical trial. Day 30 bone marrow showed complete remission. He received 3 cycles of IDAC + gilteritinib consolidation and gilteritinib maintenance with no initial plan for transplant in CR1. Relapse was documented during the $8^{th}$ month of maintenance. Salvage therapy consisted in mitoxantrone + fludarabine + cytarabine + g-csf + venetoclax resulting in CR2 and underwent haploidentical transplant 454 days after diagnosis. 5 months after transplant, molecular relapse was treated with sorafenib and 2 DLI. After which 12 months post-transplant he developed neurological disorder with *blastic meningitidis* documented on lumbar punction. He expired of progressive disease 853 days post diagnosis.

**Nancy_19 (EMD 33)**
2022 ICC Classification: **AML with inv(16)(p13.1q22) or t(16;16)(p13.1;q22)/*CBFB::MYH11***
2022 ELN risk classification: **Favorable**
25-year-old female diagnosed with AML M. Blood counts at diagnosis with WBC 6,700 with 34% circulating blasts, Hb 11.6 Platelet 148,000. Bone marrow with 59% blasts. Cytogenetic: 46,XX,inv(16)(p13q22) [19]/47,dl,+9 [5]/56,sl,+8,+10,+12,+13,+14,+19,+20,+21,+22 [6]/46,XX karyotype and FISH with CBFB/MYH11. Molecular diagnostic detected CBFb::MYH11 transcript, NGS was not performed at diagnosis. Risk category: favorable. She received induction chemotherapy with daunorubicin and cytarabine. Bone marrow aspiration at count recovery with morphologic remission. She received 3 cycles of HDAC consolidation with no initial plan for transplant in CR1. She relapsed 304 days after diagnosis and received salvage therapy with mitoxantrone + cytarabine + gemtuzumab ozogamycin resulting in CR2 and underwent matched related transplant 416 days after diagnosis. 12 months after transplant, molecular relapse was treated with 11 cycles of azacytidine 4 DLI. Meningeal syndrome justified lumbar punction revealing relapse with *central nervous system involvement* associated with marrow molecular relapse 46 months post diagnosis. She received venetoclax + multiple intrathecal injections of cytarabine + hydrocortisone, resulting in CR3 and underwent second allogenic transplant haploidentical 51 months after diagnosis. She received 5 prophylactic DLI and remains in CR3 at last follow up.

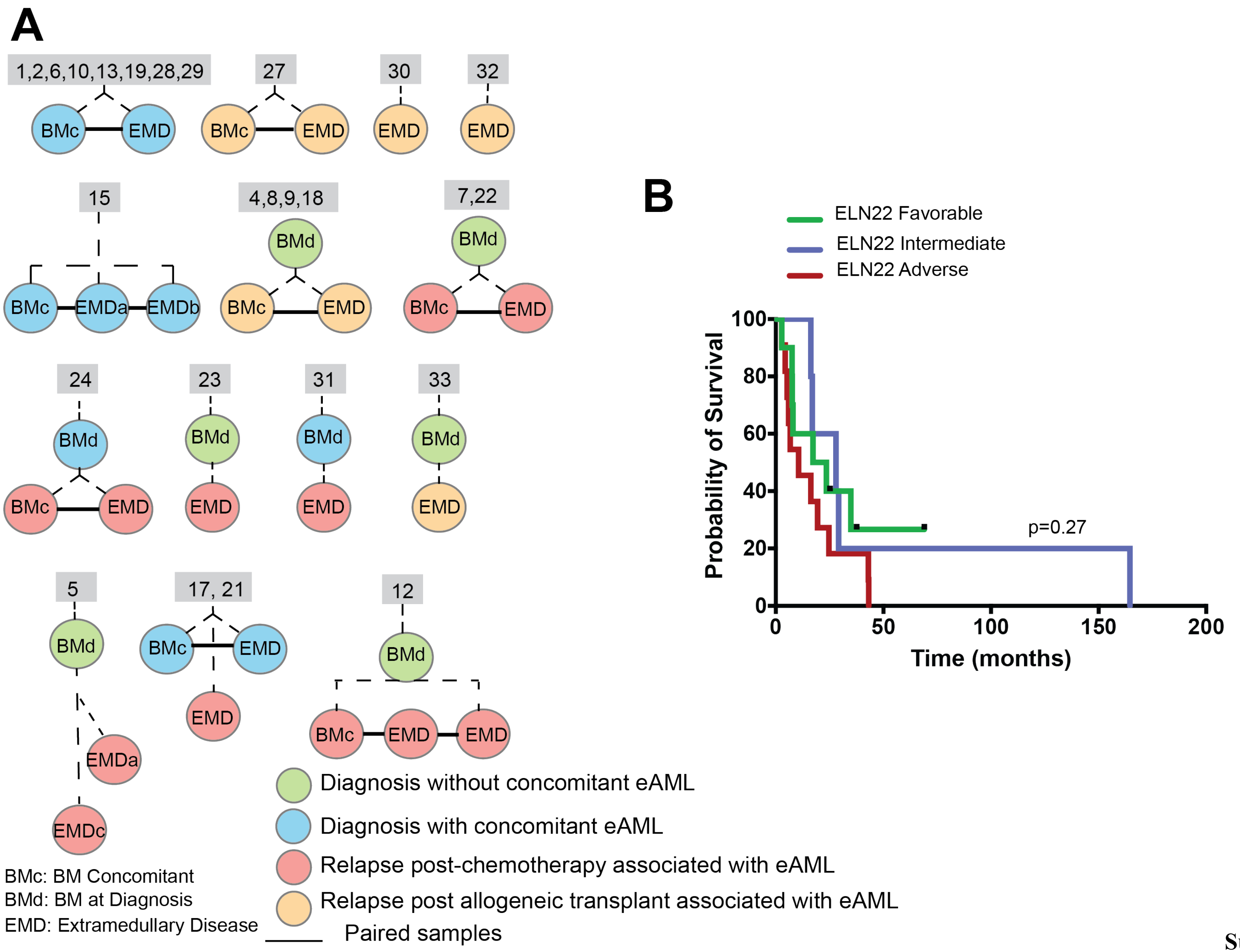


Supplementary

### Figures

**Figure S1.**

**(A) Sample timing and pairing for eAML cases.** Circles represent individual patient samples, color-coded by disease compartment: diagnosis with concomitant eAML (blue), diagnosis without concomitant eAML (green), relapse post-chemotherapy associated with eAML (red), and relapse post-allogeneic transplant associated with eAML (orange). Lines indicate paired bone marrow and extramedullary samples (obtained at the same time). Numbers within the grey squares denote patient identifiers.

**(B) Overall survival of patients stratified by ELN22 risk group.** Kaplan-Meier survival curves show the probability of overall survival over time, with risk groups color-coded by ELN22: favorable risk (green), intermediate risk (blue), and high risk (red) cases. Censored events are indicated by tick marks. The x-axis represents time in months, while the y-axis represents overall survival probability at the time of writing.

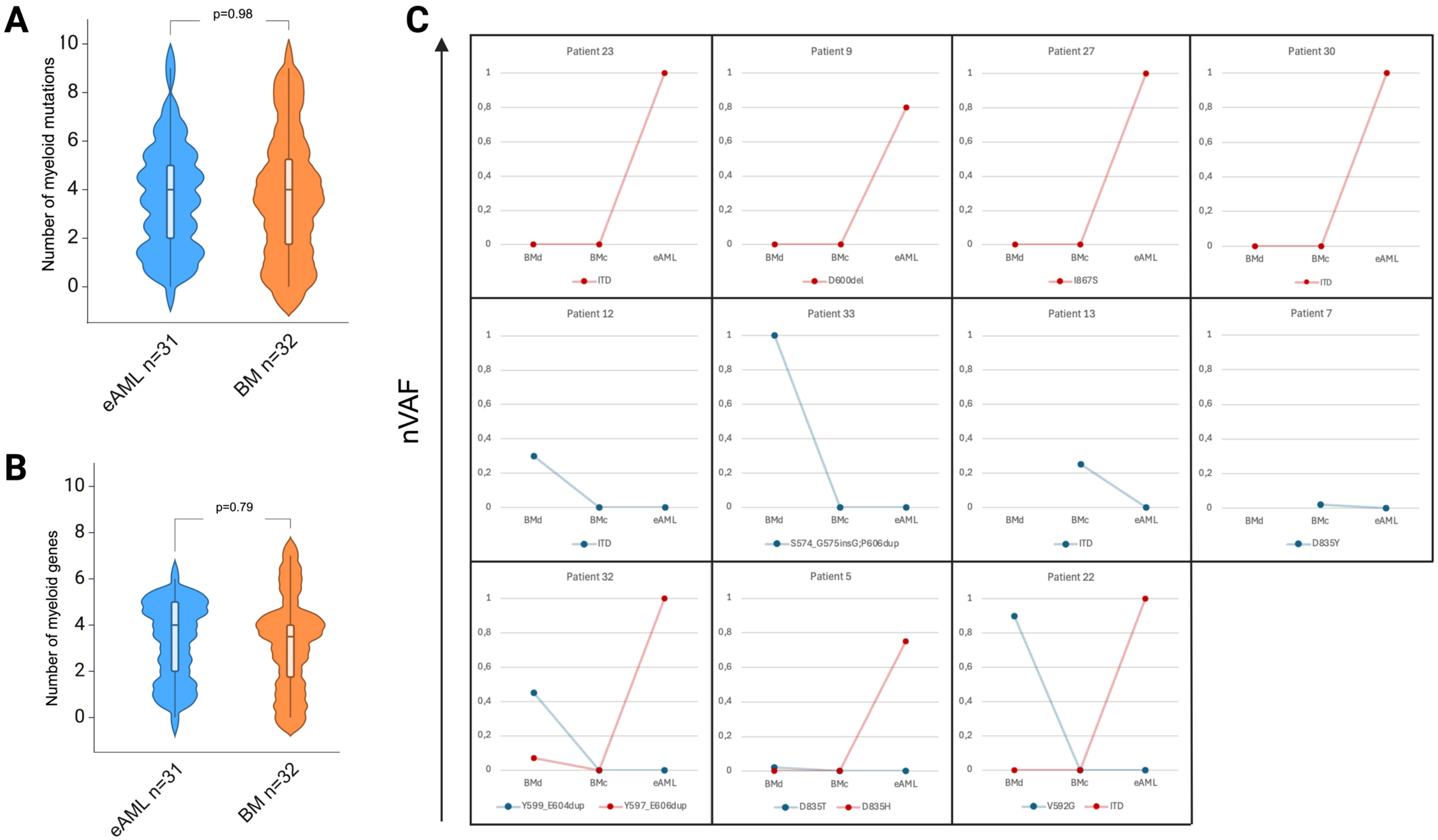
A
p=0.98
Number of myeloid mutations
10
8
6
4
2
0
eAML n=31
BM n=32
B
p=0.79
Number of myeloid genes
10
8
6
4
2
0
eAML n=31
BM n=32
C
nVAF
Patient 23
ITD
Patient 9
D600del
Patient 27
I867S
Patient 30
ITD
Patient 12
ITD
Patient 33
S574_G575insG;P606dup
Patient 13
ITD
Patient 7
D835Y
Patient 32
Y599_E604dup
Y597_E606dup
Patient 5
D835T
D835H
Patient 22
V592G
ITD
1
0,8
0,6
0,4
0,2
0
BMd
BMc
eAML

**Figure S2.**

**(A-B)** Violin plots showing the average number of mutations in myeloid genes (A) and the average number of mutated myeloid genes (B) in bone marrow samples (BM; orange) versus extramedullary AML samples (eAML; blue).

(**C**) Line plots showing the changes in normalized VAF for FLT3 mutations. Each graph represents a single patient, showing the changes in normalized variant allele frequency of FLT3 mutations in BM diagnostic samples (BMd), bone marrow samples paired to the extramedullary lesion (BMc), and extramedullary samples (eAML). Red lines indicate the emergence or expansion of mutations in eAML, whereas blue lines indicate loss or decrease in mutation frequency

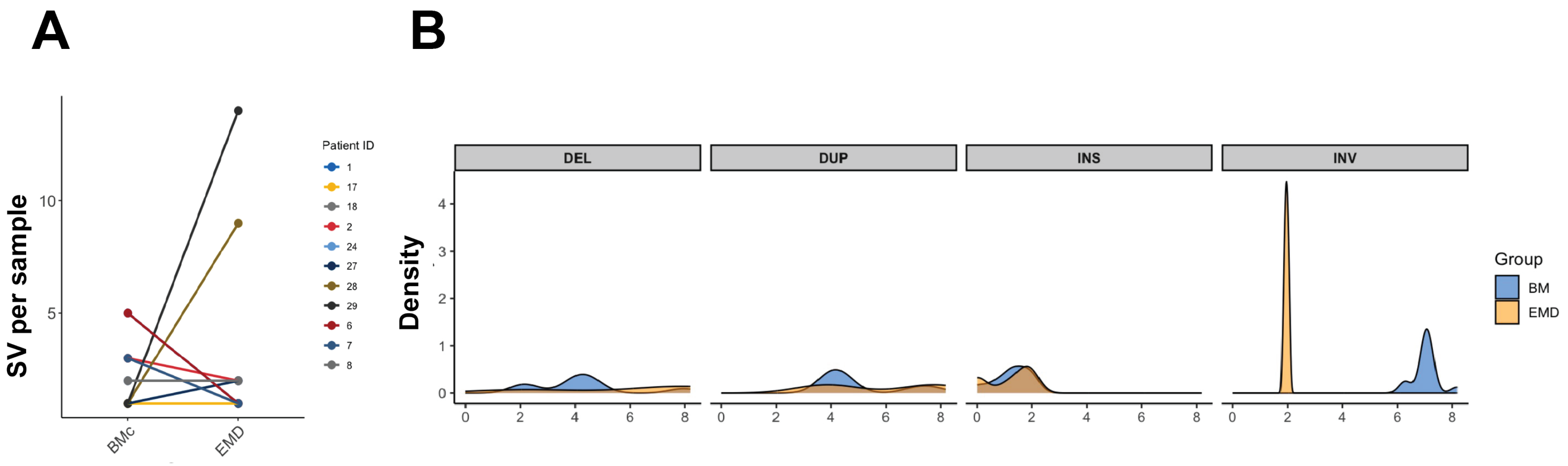


**Figure S3.**
**(A)** Line plot illustrating changes in structural variant (SV) burden between concomitant BM (BMc) and extramedullary (EMD) samples for individual patients.

**(B)** Density plot depicting the distribution of SV lengths (log10 bp) for different SV types (DEL=deletion, DUP=duplication, INS=insertion, INV=inversion) in BM (blue) and EMD (orange).

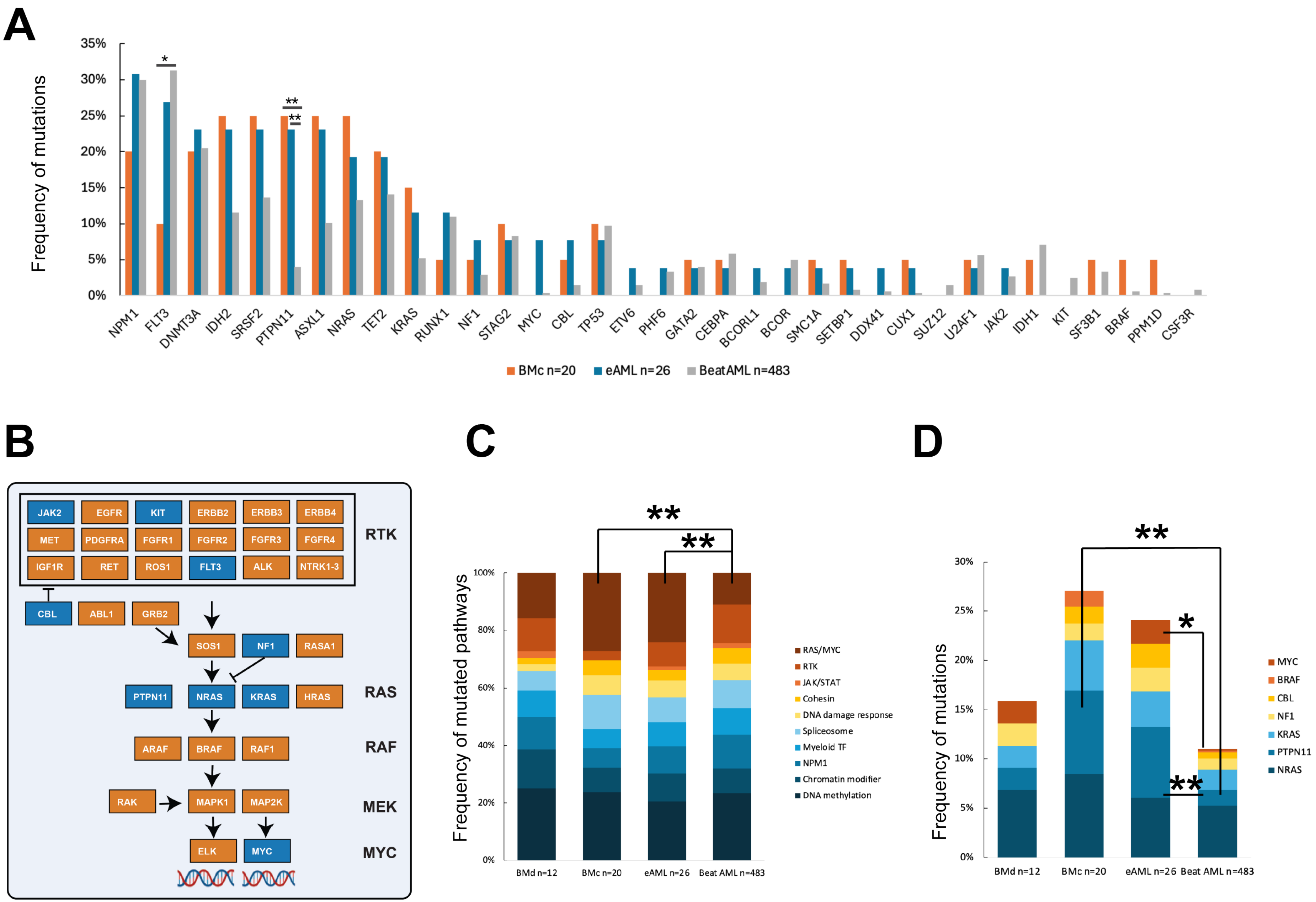
A
Frequency of mutations
35%
30%
25%
20%
15%
10%
5%
0%
*
**
**
NPM1
FLT3
DNMT3A
IDH2
SRSF2
PTPN11
ASXL1
NRAS
TET2
KRAS
RUNX1
NF1
STAG2
MYC
CBL
TP53
ETV6
PHF6
GATA2
CEBPA
BCORL1
BCOR
SMC1A
SETBP1
DDX41
CUX1
SUZ12
U2AF1
JAK2
IDH1
KIT
SF3B1
BRAF
PPM1D
CSF3R
BMc n=20
eAML n=26
BeatAML n=483
B
JAK2
EGFR
KIT
ERBB2
ERBB3
ERBB4
MET
PDGFRA
FGFR1
FGFR2
FGFR3
FGFR4
IGF1R
RET
ROS1
FLT3
ALK
NTRK1-3
RTK
CBL
ABL1
GRB2
SOS1
NF1
RASA1
PTPN11
NRAS
KRAS
HRAS
RAS
ARAF
BRAF
RAF1
RAF
RAK
MAPK1
MAP2K
MEK
ELK
MYC
MYC
C
Frequency of mutated pathways
100%
80%
60%
40%
20%
0%
**
**
BMd n=12
BMc n=20
eAML n=26
Beat AML n=483
RAS/MYC
RTK
JAK/STAT
Cohesin
DNA damage response
Spliceosome
Myeloid TF
NPM1
Chromatin modifier
DNA methylation
D
Frequency of mutations
30%
25%
20%
15%
10%
5%
0%
**
*
**
BMd n=12
BMc n=20
eAML n=26
Beat AML n=483
MYC
BRAF
CBL
NF1
KRAS
PTPN11
NRAS

**Figure S4**.

(**A**) Comparison of the frequency of myeloid gene mutations detected in bone marrow samples from patients with eAML (*n*=26) and paired BMc samples (*n*=20) to a large dataset of de novo AML cases from the BeatAML trial (*n*=483), excluding cases associated with myeloid sarcoma.

(**B**) Schematic of the receptor tyrosine kinase (RTK) and RAS pathways. In blue are genes recurrently mutated in eAML cohort.

(**C**) Distribution of mutations across functional categories in BMd (*n*=12), BMc (*n*=20), eAML (*n*=26), and BeatAML (*n*=483) cohorts. Categories include DNA methylation (dark blue), chromatin modifiers (blue), *NPM1* (light blue), myeloid transcription factors (cyan), spliceosome (light teal), DNA damage response (light yellow), cohesin complex (orange), JAK/STAT signaling (yellow), receptor tyrosine kinases (RTK) (brown), and *RAS/MYC* pathway alterations (dark brown).

**(D)** Frequency of mutations in genes related to the *RAS/MYC* pathway across BMd, BMc, eAML, and BeatAML cohorts. Genes included are *NRAS* (dark blue), *PTPN11* (blue), *KRAS* (cyan), *NF1* (yellow), *CBL* (orange), *BRAF* (light red), and *MYC* (red). Statistically significant differences are indicated by $\boldsymbol{p}<0.05$ (*) and $\boldsymbol{p}<0.01$ (**).

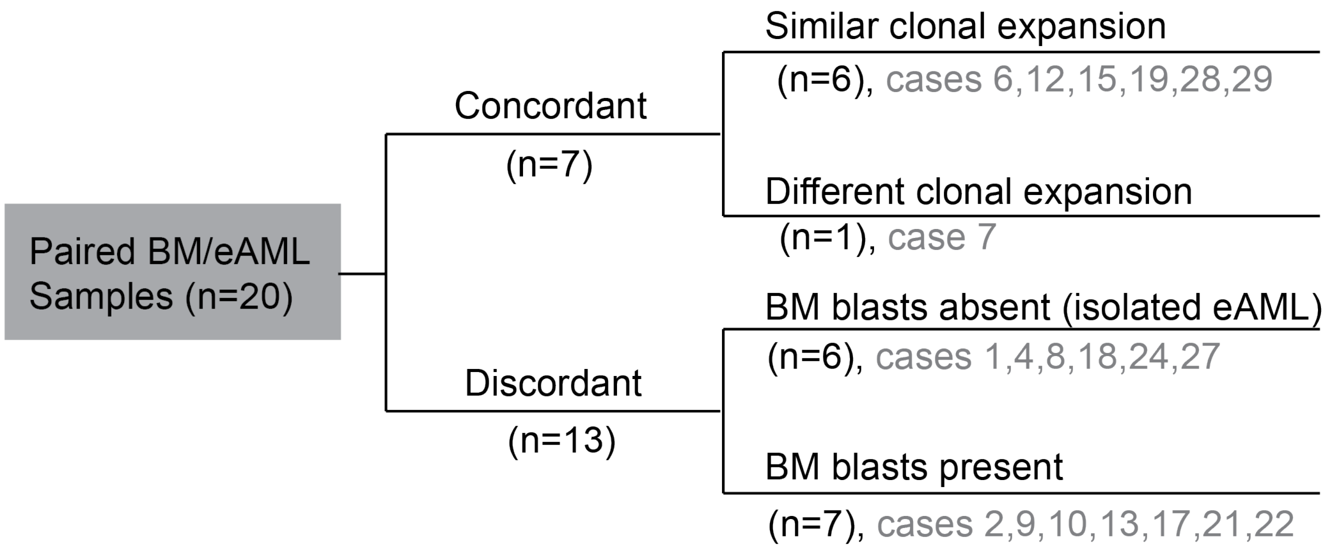


**Figure S5 .**

**Site-specific clonal evolutionary trajectories.** Categorization of the 20 paired extramedullary AML (eAML) and bone marrow (BM), providing a structured framework for understanding site-specific clonal evolution.

**Concordant mutational status (n=7):** Cases where the BM and eAML shared highly similar mutational profiles, suggesting a common clonal origin with minimal divergence. Nearly all of these cases exhibit equivalent variant allele frequency (nVAF) of key mutations in BM and eAML, indicating parallel evolution of dominant clones. Only one case exhibited different nVAF between BM and eAML, suggesting selection pressures leading to site-specific subclonal dominance (Patient IDs: 7)

**Discordant mutational status (n=14):** Cases where BM and eAML have distinct mutational landscapes, indicating independent clonal evolution. These cases are further classified into: Concomitant BM Involvement (n=7): cases with leukemia detected in both BM and extramedullary sites. Isolated eAML (n = 6All these cases exhibit discordant mutational profiles compared to BM.

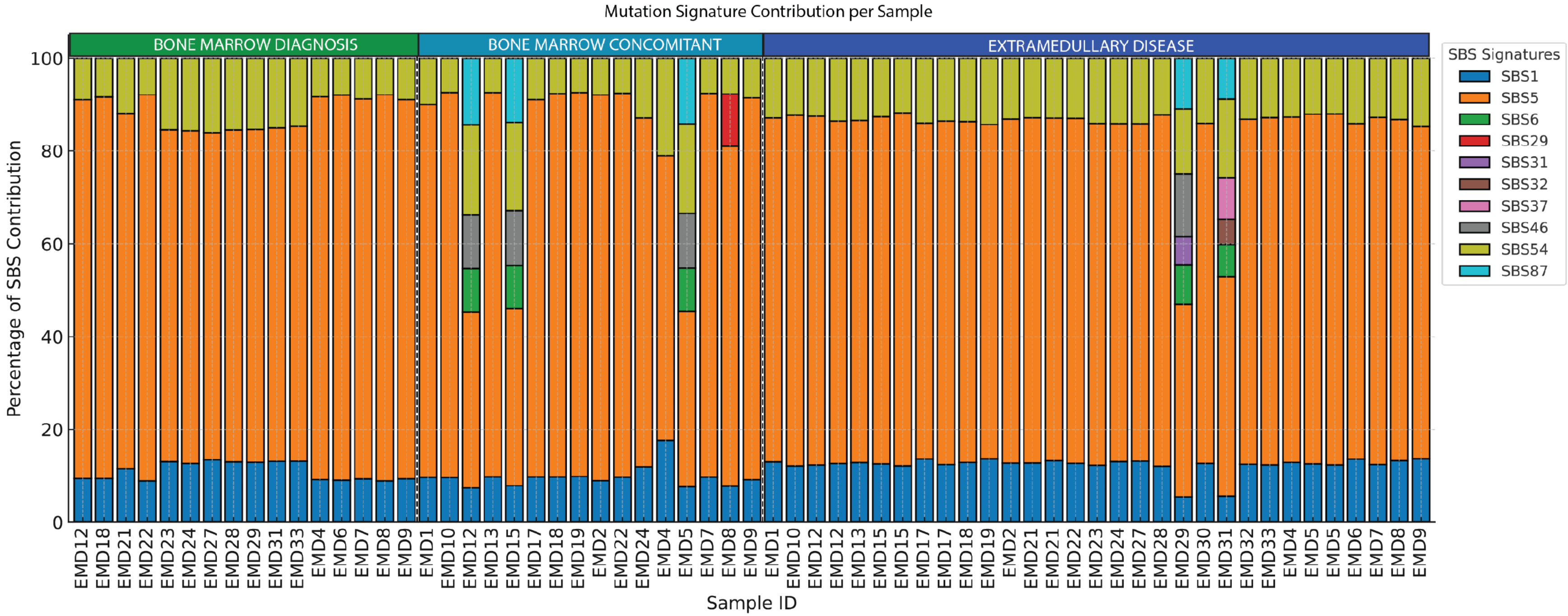


**Figure S6:** Stacked bar plot illustrating the contribution of COSMIC single-base substitution (SBS) mutational signatures across three conditions – Bone Marrow at Diagnosis, Bone Marrow Concomitant, and Extramedullary Disease. Each vertical bar represents an individual AML sample, with different colored segments denoting the fraction of mutations attributed to distinct SBS mutational signatures.

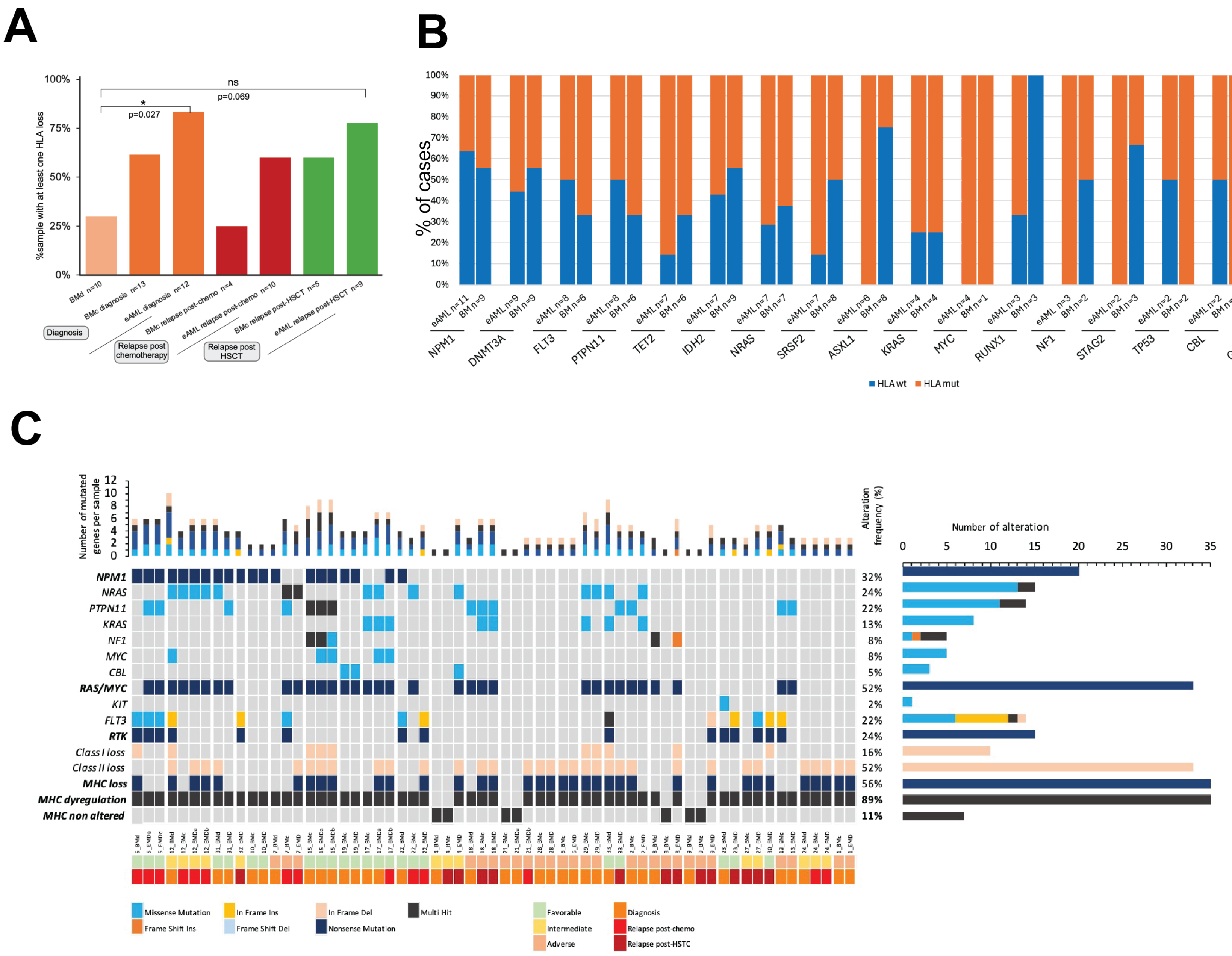
A
100%
75%
50%
25%
0%
%sample with at least one HLA loss
ns
p=0.069
*
p=0.027
BMd n=10
BMc diagnosis n=13
eAML diagnosis n=12
BMc relapse post-chemo n=4
eAML relapse post-chemo n=10
BMc relapse post-HSCT n=5
eAML relapse post-HSCT n=9
Diagnosis
Relapse post chemotherapy
Relapse post HSCT
B
% of cases
NPM1
DNMT3A
FLT3
PTPN11
TET2
IDH2
NRAS
SRSF2
ASXL1
KRAS
MYC
RUNX1
NF1
STAG2
TP53
CBL
GATA2
CEBPA
BCORL1
HLA wt
HLA mut
C
Number of mutated genes per sample
NPM1
NRAS
PTPN11
KRAS
NF1
MYC
CBL
RAS/MYC
KIT
FLT3
RTK
Class I loss
Class II loss
MHC loss
MHC dyregulation
MHC non altered
Alteration frequency (%)
32%
24%
22%
13%
8%
8%
5%
52%
2%
22%
24%
16%
52%
56%
89%
11%
Number of alteration
Missense Mutation
Frame Shift Ins
In Frame Ins
Frame Shift Del
In Frame Del
Nonsense Mutation
Multi Hit
Favorable
Intermediate
Adverse
Diagnosis
Relapse post-chemo
Relapse post-HSTC

**Figure S7.**

(**A**) Frequency of HLA altered status among sample group and disease course status: BM at diagnosis without associated eAML (n=10), BM concomitant to eAML at diagnosis (n=12), extramedullary lesion at diagnosis (n=11), BM concomitant to eAML at relapse post-chemotherapy (n=4), extramedullary lesion at relapse post-chemotherapy (n=9), BM concomitant to eAML at relapse post-HSTC (n=6),extramedullary lesion at relapse post-HSCT (n=11). Statistically significant differences are indicated by $p<0.05$(*). HSTC: hematopoietic stem cell transplantation

(**B**) Distribution of wild type (blue) or altered (orange) HLA status among RMGs with at least 3 mutations within the 63 samples.; RMG: recurrently mutated gene in AML

(**C**) Oncoplot showing the most recurrently mutated pathways (in bold), highlighting the genetic alterations associated with MHC dysregulation in AML. Each column represents an individual AML case, and each row corresponds to a specific genetic event in the pathway/gene listed. Mutation types are color-coded: missense mutations (blue), frameshift insertions (orange), inframe insertions (yellow), frameshift deletions (light blue), inframe deletions (light orange), nonsense mutations (dark blue), and multi-hit mutations (black). The annotation bar at the bottom indicates the clinical status of each patient: ELN22 favorable (light green), ELN22 intermediate (yellow), ELN22 adverse (light orange), diagnosis (orange), relapse post-chemotherapy (red), and relapse post-hematopoietic stem cell transplantation (dark red). The right panel displays the frequency of mutations per gene/pathway.
